# The final step of 40S ribosomal subunit maturation is controlled by a dual key lock

**DOI:** 10.1101/2020.07.29.226936

**Authors:** Laura Plassart, Ramtin Shayan, Christian Montellese, Dana Rinaldi, Natacha Larburu, Carole Pichereaux, Simon Lebaron, Marie-Françoise O’Donohue, Ulrike Kutay, Julien Marcoux, Pierre-Emmanuel Gleizes, Célia Plisson-Chastang

**Affiliations:** Laboratoire de Biologie Moléculaire Eucaryote, Centre de Biologie Intégrative, Université de Toulouse, CNRS, UPS, 118 route de Narbonne, 31062 Toulouse Cedex, France; Institut für Biochemie, ETH Zürich, 8093 Zurich, Switzerland; Institut de Pharmacologie et Biologie Structurale, Université de Toulouse, CNRS, UPS, 205 route de Narbonne, 31062 Toulouse Cedex, France; Institute of Structural and Molecular Biology, Birkbeck, University of London, Malet Street, WC1E7 London, UK; CSL Behring, CSL Biologics Research Center, Freiburgstrasse 3, 3010 Bern, Switzerland; Department of Life Sciences, Sir Ernst Chain Building, Imperial College London, SW72AZ London, UK

## Abstract

Preventing premature interaction of preribosomes with the translation apparatus is essential to translation accuracy. Hence, the final maturation step releasing functional 40S ribosomal subunits, namely processing of the 18S ribosomal RNA 3’ end, is safeguarded by protein DIM2, which both interacts with the endoribonuclease NOB1 and masks the rRNA cleavage site. To elucidate the control mechanism that unlocks NOB1 activity, we performed cryo-EM analysis of late human pre-40S particles purified using a catalytically-inactive form of ATPase RIO1. These structures, together with in vivo and in vitro functional analyses, support a model in which ATP-loaded RIO1 cooperates with ribosomal protein RPS26/eS26 to displace DIM2 from the 18S rRNA 3’ end, thereby triggering final cleavage by NOB1; release of ADP then leads to RIO1 dissociation from the 40S subunit. This dual key lock mechanism requiring RIO1 and RPS26 guarantees the precise timing of pre-40S particle conversion into translation-competent ribosomal subunits.

## INTRODUCTION

Synthesis of eukaryotic ribosomes relies on a large array of ribosome biogenesis factors (RBFs) that coordinate the multiple steps of pre-ribosomal RNA (pre-rRNA) modification, cleavage and folding, together with ribosomal protein (RP) assembly (Bohnsak and Bohnsack, 2019). Progression through this process, defined by the timely association or dissociation of RBFs and RPs to pre-ribosomal particles and the gradual maturation of pre-rRNAs, is tightly monitored from one stage to the next. These monitoring mechanisms not only ensure quality control along this intricate biosynthetic pathway, but also prevent binding of immature ribosomal subunit precursors (pre-ribosomes) to mRNAs or to components of the translation apparatus. Such interactions would not only interfere with ribosome biogenesis, but also affect translation accuracy. The recent discovery of congenital diseases or cancers linked to ribosome biogenesis defects and translation dysregulation underscores the importance of the control mechanisms that license newly formed ribosomal subunits to enter the translation-competent pool of ribosomes (Aubert et al., 2018; Bohnsak and Bohnsack, 2019; Sulima et al., 2017).

After initial assembly in the nucleolus, pre-40S particles are rapidly transported to the cytoplasm (Rouquette et al., 2005), where translation takes place. Cytoplasmic precursors to the 40S ribosomal subunits (pre-40S particles) closely resemble their mature counterparts (Ameismeier et al., 2018; Larburu et al., 2016), which possess binding sites for numerous components of the translation machinery including components of the translation initiation complex, tRNAs and mRNAs. Thus, pre-40S particles would be especially prone to premature interactions with the translation apparatus without the presence of several RBFs (Bystin/ENP1, LTV1, RIO2, TSR1, DIM2, NOB1), that occupy the binding sites of translation partners near their “head” and “platform” domains (Larburu et al., 2016; Strunk et al., 2012). These two structural domains undergo several remodeling steps in the cytoplasm leading to the gradual release of these RBFs (Ameismeier et al., 2018; Zemp et al., 2009; Zemp et al., 2014). This maturation process ends with the cleavage of the 18S rRNA 3’ end and the dissociation of the last RBFs, DIM2 and NOB1, which converts pre-40S particles into functional subunits. Interfering with this late stage can result in the incorporation of immature 40S subunits in the translation pool (Belhabich-Baumas et al., 2017; Parker et al., 2019).

Up to this final stage, the 3’ end of the 18S rRNA is extended with remnants of the internal transcribed spacer 1 (ITS1). In human cells, this last precursor to the 18S rRNA (called 18S-E pre-rRNA) is generated in the nucleolus by endonucleolytic cleavage of earlier pre-rRNAs 78 or 81 nucleotides downstream of the mature 3’ end (Preti et al., 2013; Sloan et al. 2013; Tafforeau et al. 2013). This 3’ tail is then gradually trimmed by exonucleases before and after nuclear export, including PARN in the nucleus (Montellese et al., 2017). However, processing of the 18S rRNA 3’ end is finalized by endonuclease NOB1, which cleaves at the so-called site 3. NOB1 is already incorporated into the pre-40S particle in the nucleolus on the platform domain near site 3, but it is restricted from cleaving the rRNA by DIM2/PNO1, another RBF, which contacts NOB1 in the particle and masks the cleavage site on the RNA (Ameismeier et al., 2018; Larburu et al., 2016). This conformation also maintains a large gap between the catalytic site of NOB1 and its substrate. Dissociation of NOB1 and DIM2 from the pre-40S particles only occurs with cleavage of the 18S rRNA 3’ end. In addition, NOB1 lays in the mRNA binding cleft and prevents association of pre-40S particles with mRNAs until this ultimate maturation step (Parker et al., 2019). Thus, NOB1 and DIM2 constitute a critical checkpoint controlling the release of nascent 40S subunits into the translating pool.

The molecular mechanism driving the accurate activation of rRNA cleavage by NOB1 remains poorly known, but the RIO1 ATPase was shown to play a critical function in this process (Widmann et al., 2012). RIO1 briefly associates with pre-40S particles before the final cleavage step. Like RIO2, another ATPase of the same family that intervenes earlier in pre-40S particle maturation, RIO1 adopts different conformations depending on its nucleotide binding state (Ferreira-Cerca et al., 2014). Absence of RIO1 or suppression of its catalytic activity impairs both rRNA cleavage and release of NOB1 and DIM2 (Ferreira-Cerca et al., 2014; Widmann et al., 2012). In addition, incorporation into pre-40S particles of the last ribosomal protein, RPS26/eS26, takes place at this stage, in contact with the 18S rRNA 3’ end. RPS26 was shown to be necessary for efficient cleavage by NOB1 (O’Donohue et al., 2010), but its interplay with RIO1, DIM2 and NOB1 remains unclear.

Here, we have used cryo-electron microscopy (cryo-EM) to solve the structures of human pre-40S particles trapped with a catalytically inactive form of RIO1 in this final maturation stage. Image analysis revealed two distinct structural states, prior and after 18S rRNA cleavage. In the pre-cleavage state, DIM2 and NOB1 are still in place, while RIO1 position is not clearly defined and RPS26 is absent. In contrast, the post-cleavage state displays RIO1 and RPS26 stably associated to 40S particles, which contain mature 18S rRNA. In vivo assays confirmed the central role of RPS26 for triggering both processing of the the 18S-E pre-rRNA processing and the release of DIM2, NOB1 and RIO1. In vitro, cleavage of the 18S-E pre-rRNA was partially stimulated by ATP binding to RIO1. These data suggest a model, in which ATP-bound RIO1 and RPS26 cooperatively displace DIM2 to activate the final cleavage of the 18S rRNA 3’ end by NOB1.

## RESULTS

### Pre-40S particles purified with a catalytically-deficient form of RIO1 display pre- and post-18S rRNA processing structural states

In order to characterize the final cytoplasmic maturation steps undergone by human pre-40S particles, we purified pre-40S particles from a human cell line overexpressing a tagged version of a catalytically-inactive form of RIO1 mutated on aspartic acid 324 (D324A) (Widmann et al., 2012). Autophosphorylation of this mutant in the presence of ATP was previously shown to be strongly impaired (Widmann et al., 2012). Protein and RNA composition of the affinity-purified complexes, hereafter called RIO1(kd)-StHA pre-40S particles, were analyzed using SDS-PAGE, Northern blot and bottom-up proteomics (Figure 1). SDS-PAGE revealed a complex protein pattern with a typical trail of low molecular weight bands corresponding to ribosomal proteins. Northern blot showed that the 18S-E pre-rRNA was the only 18S rRNA precursor precipitated by this bait, while no rRNA of the large ribosomal subunit could be detected (Figure 1, figure supplement 1). Bottom-up proteomics confirmed that the catalytically-dead version of RIO1 associates with the methylosome, a complex composed of proteins PRMT5 and MEP50 that methylates proteins involved in gene expression regulation (Guderian et al., 2011), in addition to late pre-40S particles (Montellese et al., 2020; Widmann et al., 2012). Of note, these purified pre-40S particles contain all ribosomal proteins of the small subunit (RPSs), including the late binding RPS10 and RPS26, but only a handful of ribosome biogenesis factors, namely RIO1 (the bait), DIM2, NOB1 and TSR1 (Figure 1 and figure supplement 2). Other proteins also copurified with RIO1(kd)-StHA: while some can unambiguously be identified as components of the translation apparatus (eIF1AD, eIF4A, RPL1, …), it is not known whether others (casein kinases CK2*α*1 and CK2*α*2, …) belong to the methylosome, ribosomal particles or yet another co-purified complex.

**Figure 1.**
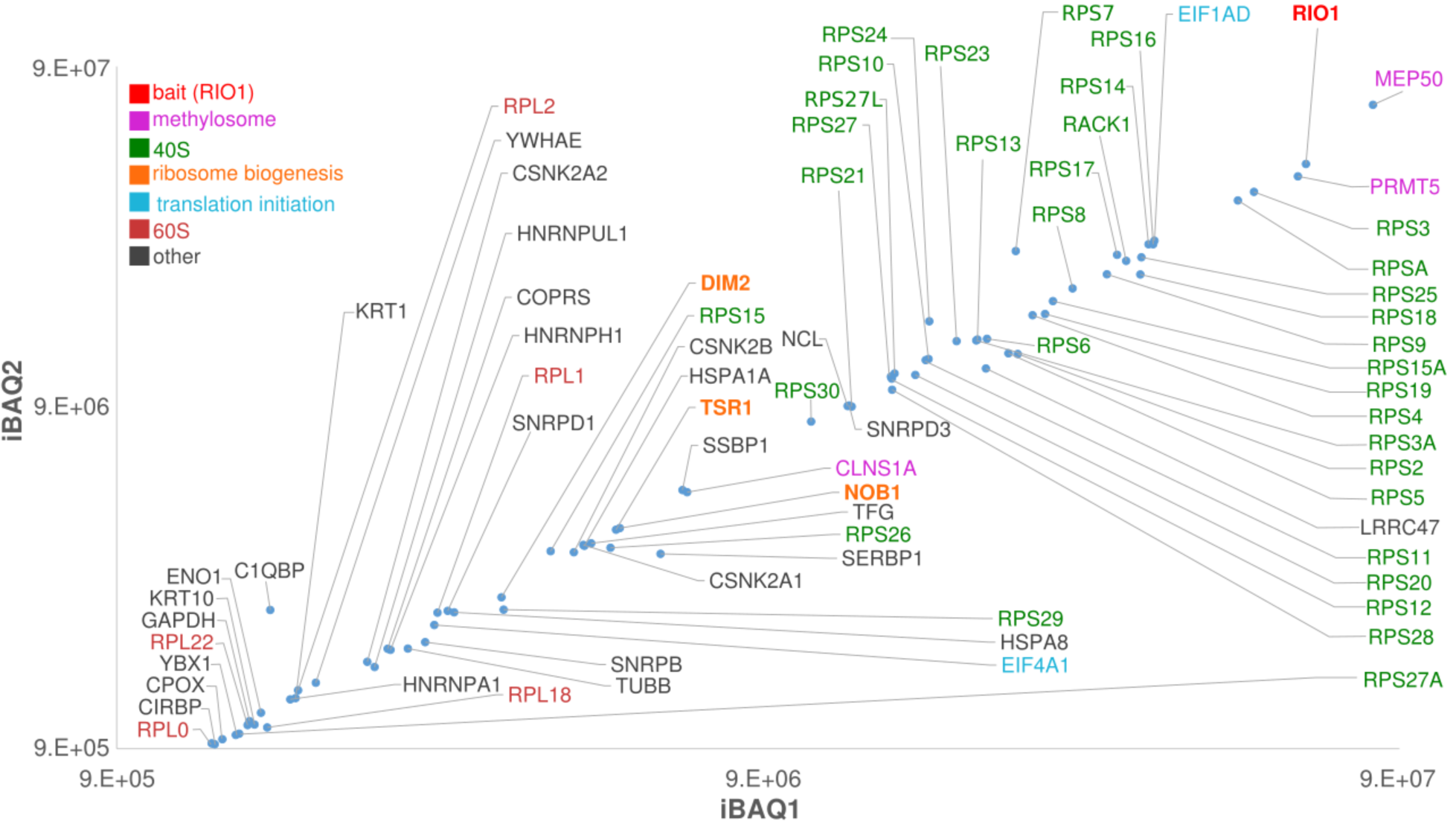
Label-free bottom-up proteomic analysis of RIO1(kd)-StHA co-purified proteins. The most intense proteins (first two logs) with an observed/observable peptide ratio > 30% are displayed and color coded as indicated on the graph. Three independent experimental replicates were performed. The current plot represents iBAQs (intensity-Based Absolute Quantification) of experimental replicate 1 (iBAQ1) against experimental replicate 2 (iBAQ2). The iBAQ value is obtained by dividing protein intensities by the number of theoretically observable tryptic peptides (Schwanhäusser et al., 2011). Other plots are displayed in Figure1 - figure supplement 1. Of note USP16, which was recently identified as a key player in both ribosome biogenesis and translation (Montellese et al., 2020) was also identified among RIO1(kd)-StHA proteic partners. It ranked at the 62th position when quantifying proteins based on their normalized abundances, and at the 98th position when quantifying proteins based on their iBAQs, ie. normalized abundance divided by the number of theoretically observable peptides (see the PRIDE repository (Perez-Riverol et al., 2019); dataset identifier: PXD019270).

We then performed cryo-EM and single particle analysis on the particles purified using RIO1(kd)-StHA as bait. 2D classification assays performed with RELION (Scheres, 2012) yielded class sums corresponding to pre-40S particles as well as various views of the methylosome (Timm et al., 2018) (Figure 2, and Figure 2 - figure supplement 1). Views were selected for further processing; an extensive 3D classification scheme resulted in two distinct 3D structures, hereafter called state A and state B, that likely reflect two successive pre-40S maturation steps (Figure 2 and Figure 2 - figure supplement 1). Both structures, which represented ∼21 % (state A) and ∼57 % (state B) of the pre-40S particles comprised within the analysis were refined to overall resolutions of 3.2 and 3.0 Å, respectively, according to RELION’s gold standard FSC (Figure 2 - figure supplements 1 and 2).

**Figure 2.**
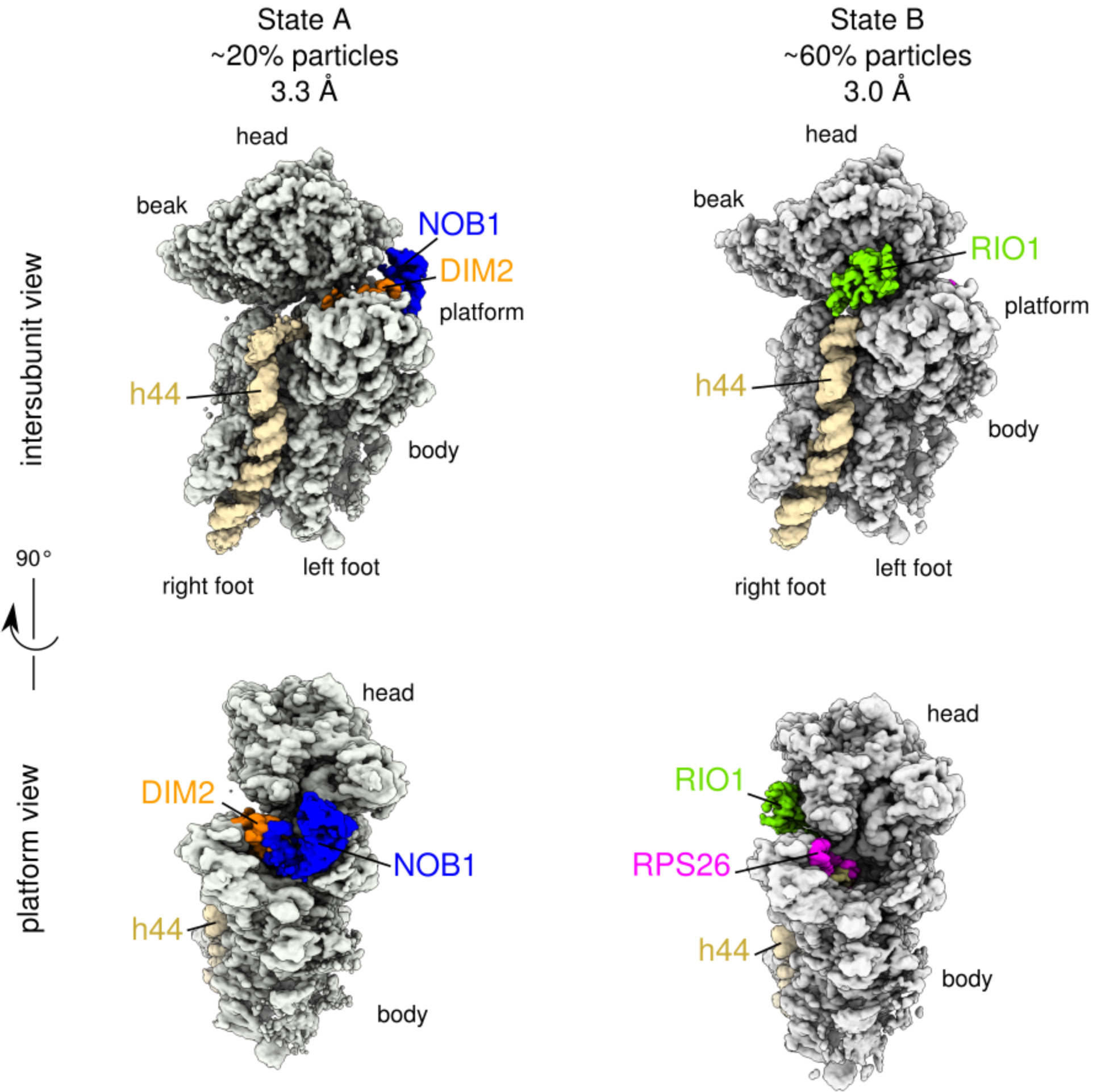
Cryo-EM and single particle analysis reveal two distinct structural states, pre- (State A) and post- (State B) 18S-E rRNA cleavage. Surface views of cryo-EM maps of RIO1(kd)-StHA pre-40S particles in structural state A (left panel) and state B (right panel). Ribosomal proteins, rRNA segments and RBFs of interest have been segmented and colored as indicated on the figure. Image processing details are shown in Figure 2 – figure supplements 1 and 2).

State A harbored an rRNA scaffold closely resembling that of the mature 18S rRNA. On the head region, the 3-way junction formed by rRNA helices h34, h35 and h38 is fully formed (Mohan et al., 2014), while RPS3, RPS10 and RPS12 occupy their final position. Like in many other pre-40S structures (Ameismeier et al., 2018; Heuer et al., 2017; Mitterer et al., 2019; Scaiola et al., 2018), the upper part of rRNA h44, located on the intersubunit side, appears detached from the body (Figure 3a). Low-pass filtering of the cryo-EM map of state A also revealed the presence of TSR1 in this region (Figure 3 - figure supplement 1). Combined with the bottom-up proteomic analysis that revealed the presence of this RBF among the co-purified proteins, this suggests that TSR1 might still be loosely bound to pre-40S particles at this maturation state. On the platform region, the cryo-EM density map allowed to unambiguously position nucleotide A1870, belonging to the ITS1 after the 3’ end of the mature 18S rRNA. In this state, DIM2 protects this region of the 18S-E pre-rRNA and impedes endonucleolytic cleavage by NOB1, located right next to DIM2 (Figure 2 and 3b). These results suggest that RIO1(kd)-StHA pre-40S particles in structural state A are in an immature state in which the rRNA scaffold harbors a quasi-mature conformation, but the remaining nucleotides of the ITS1 have not yet been cleaved off by NOB1. Furthermore, NOB1 and DIM2 are the only two RBFs that can clearly be distinguished, while RIO1(kd)-StHA, the protein used as purification bait, cannot be positioned on this cryo-EM map. This suggests that, like TSR1, RIO1(kd)-StHA is only loosely bound to pre-40S particles found in structural state A.

**Figure 3.**
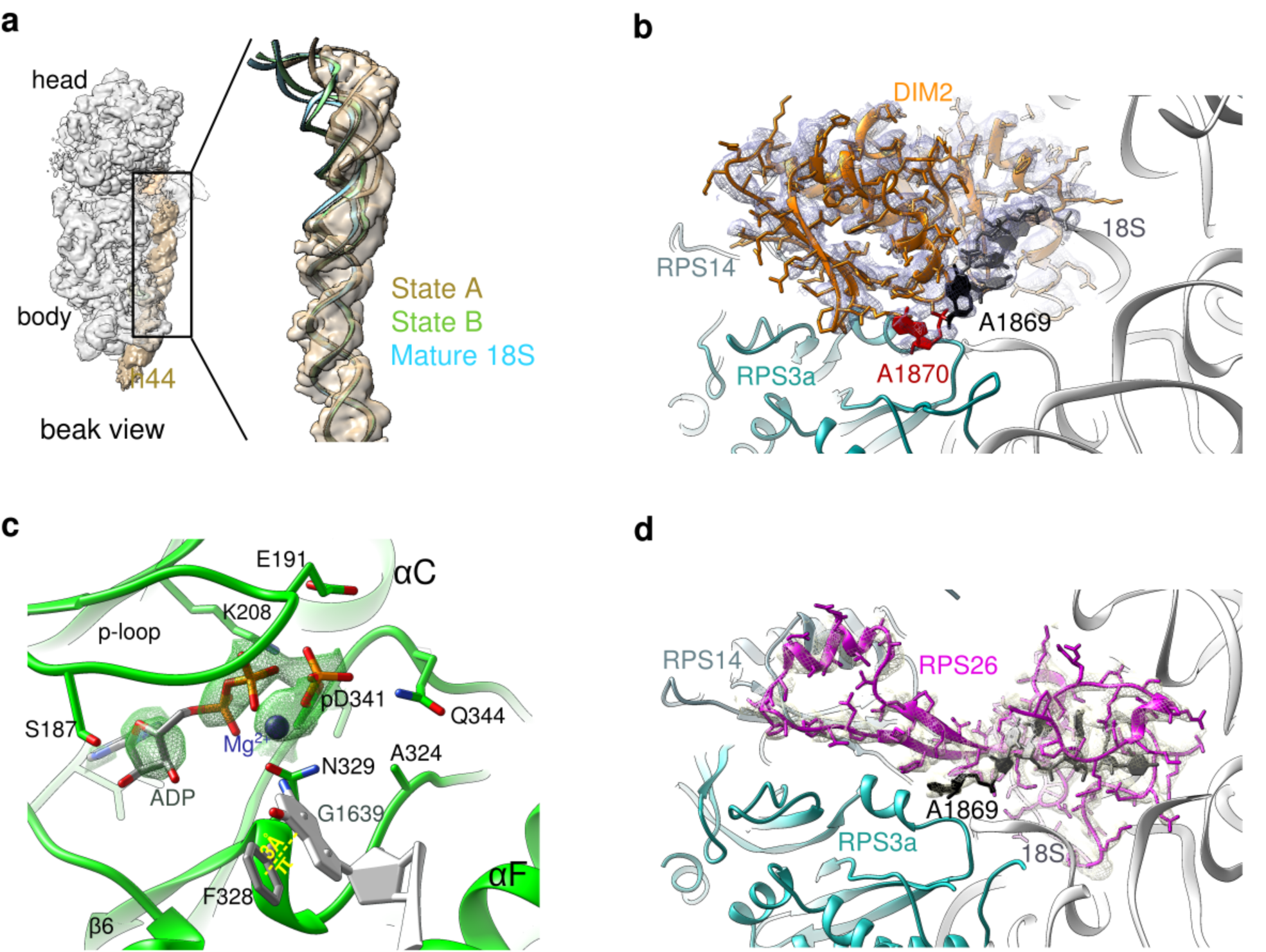
Structural details of RIO(kd)-StHA pre-40S particles. **a**, The upper part of 18S rRNA helix h44 is in an immature position in structural state A. The cryo-EM density map corresponding to this helix has been segmented (beige density); the atomic model of rRNA h44 in structural state A is represented in golden; superimposed 18S rRNA h44 as found in structural state B and in the mature 40S subunit (PDB 6EK0 (Natchiar et al., 2018)) are in green and blue, respectively. **b**, Close-up on the platform domain of structural state A. Segmented cryo-EM densities corresponding to DIM2 and 18S-E pre-rRNA are shown as a grey mesh. The 3’-end of the mature 18S rRNA (nucleotides 1865-1869) is shown in black, while A1870 in the ITS1 is in red. The 18S rRNA is otherwise shown as a grey ribbon. DIM2 is in orange, RPS3a in turquoise. NOB1 was removed from this representation for the sake of clarity. **c**, The catalytic pocket of RIO1 is in an “active” state within structural state B pre-40S particles, and carries an ADP and a phospho-aspartate (pD341). The cryo-EM density corresponding to ADP, p-Asp and Magnesium is shown as a green mesh. RIO1 is shown in green; ADP in dark grey, Mg^2+^ in dark violet, and 18S rRNA G1639 closing RIO1 catalytic domain by a pi-stacking interaction (yellow dashed line) with RIO1 Phe 328 in light grey. **d**, Close-up on the platform domain of structural state B. Segmented cryo-EM densities corresponding to RPS26 and 18S rRNA are shown as a light grey mesh. 18S rRNA 3’-end (nucleotides 1865-1869) is shown in black, while otherwise as grey ribbon. RPS26 is in magenta and RPS3a in turquoise.

On the contrary, the cryo-EM map corresponding to state B harbored a density located on the intersubunit side, at the back of the head region, in which the X-Ray structure of RIO1 could unambiguously be fitted (Ferreira-Cerca et al., 2014). Of note, RIO1 occupies a position strongly overlapping with that of RIO2 in earlier pre-40S maturation states (Ameismeier et al., 2018) (Figure 3 - figure supplement 1). This observation explains why both ATPases are not found together in pre-40S particles, and confirms that RIO1 replaces RIO2 at the back of the head (Ameismeier et al., 2018; Knüppel et al., 2018; Widmann et al., 2012). The C-terminal domain of RIO2 was shown to be deeply inserted within the body of human pre-40S particles (Ameismeier et al., 2018). In contrast, the C-terminal domain of RIO1 is not resolved here; furthermore, the upper part of rRNA helix h44 occupies its mature position, which would preclude insertion of the C-terminal domain of RIO1 at the same position as the C-terminal domain of RIO2. Indeed, the strong sequence divergence of the C-termini of RIO proteins suggests that these domains play a major role in the functional specificity of these proteins, which is supported by the structural difference observed here.

As previously observed in the structure of wild-type RIO1 solved by X-ray crystallography (Ferreira-Cerca et al., 2014), our cryo-EM map revealed that the catalytic pocket of mutant RIO1 encloses an ADP together with phospho-aspartate pD341 (Figure 3c). Thus, the D324A RIO1 mutation does not fully prevent ATP hydrolysis, but rather blocks the release of its reaction products. ATP hydrolysis is thought to be accompanied by a significant conformational change of RIO1 (Knüppel et al., 2018; Kühlbrandt, 2004), which might be essential for stable association of RIO1 to pre-40S particles. Blockade of RIO1 in this conformation is likely to explain why this point mutation traps the protein on the particles. Furthermore, the RIO1(kd) catalytic pocket appears to be closed through a pi-stacking interaction between phenylalanine F328 and 18S rRNA G1639 (Figure 3c). This highly conserved nucleotide plays a crucial role in tRNA translocation, which puts RIO1 in a good position to probe this mechanism (see discussion below).

In state B, the rRNA scaffold harbors a fully mature conformation. Contrary to what was observed on the platform in state A, no density corresponding to nucleotides belonging to the ITS1 could be detected, suggesting that the 18S rRNA 3’ end is mature. Furthermore, neither NOB1 nor DIM2 could be found in this area; instead, a cryo-EM density sheathing the 18S rRNA 3’ end was unambiguously attributed to the presence of RPS26 (Figure 3d). These observations indicate that state B corresponds to particles after rRNA cleavage by NOB1. This conclusion was further supported by the analysis of the 3’-end of the 18S rRNA by RNase H digestion assays (Figure 4a). Early and intermediate cytoplasmic pre-40S particles purified using tagged forms of LTV1 or DIM2 as baits (Ameismeier et al., 2018; Wyler et al., 2011) contained a large majority of 18S-E pre-rRNA relative to 18S rRNA. Most of the 18S-E rRNA in these particles included 4 to 9 nucleotides of the ITS1 (Figure 4 and Figure 4-Figure supplement 1). In stark contrast, the 18S rRNA was four times more abundant than the 18S-E precursor in RIO1(kd)-StHA pre-40S particles, thus confirming that a majority of these particles contain mature 18S rRNA (Figure 4, b and c).

**Figure 4.**
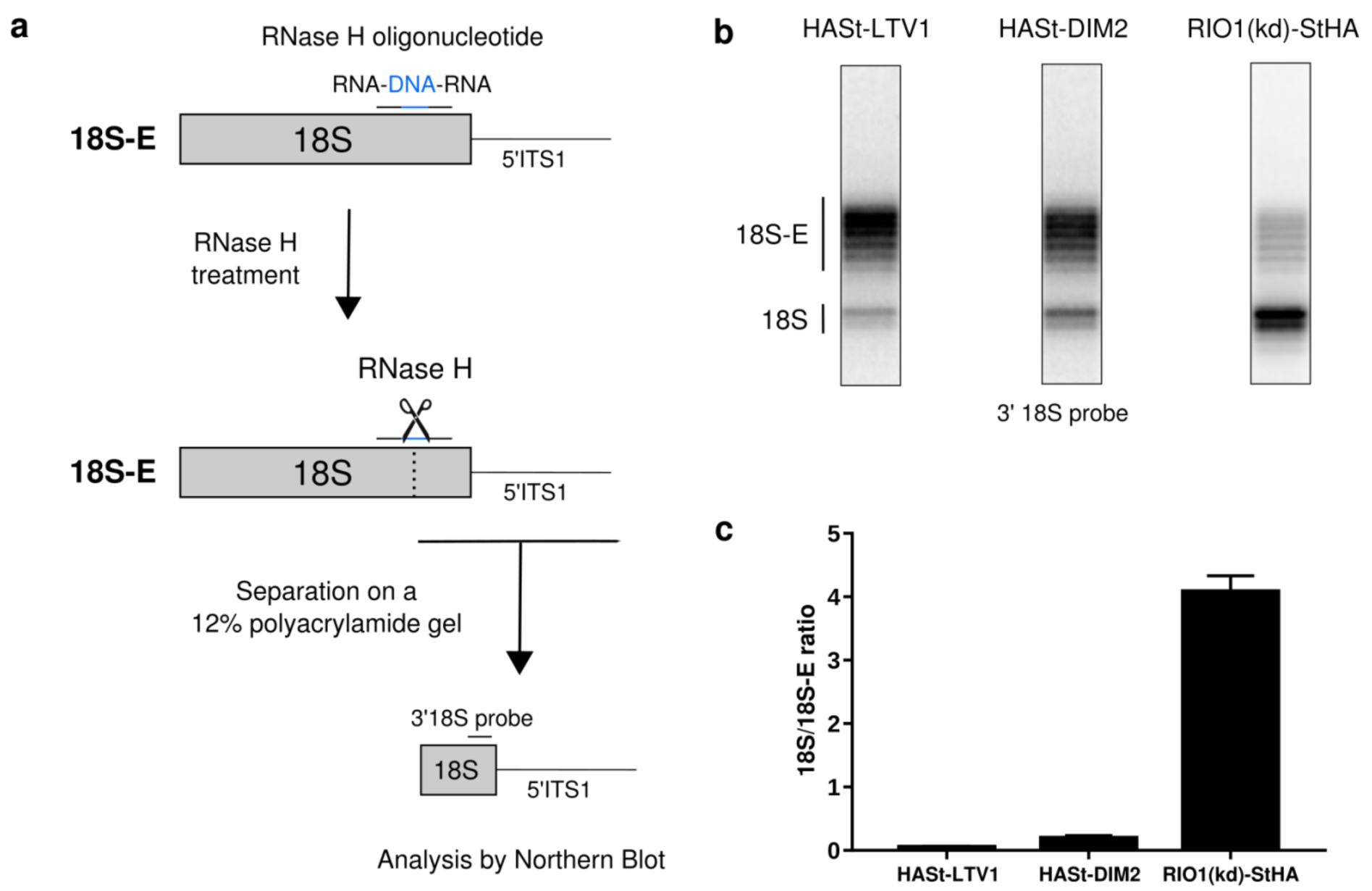
Late RIO1(kd)StHA pre-40S particles contain a high proportion of mature 18S rRNA. **a**, Diagram representing steps of the pre-40S rRNA digestion by RNase H. **b**, RNase H assays were performed on RNAs extracted from pre-40S particles purified with the mentioned StHA tagged bait, and separated on a 12% polyacrylamide gel. The 18S rRNA and its precursors were revealed by the 3’18S radiolabeled probe. Bands are separated with single nucleotide resolution, as shown in Figure 4 – figure supplement 1. **c**, Signals corresponding to the 18S-E and 18S rRNAs were quantified by phosphorimaging and represented by the 18S/18S-E ratio for the different purified pre-40S particles. The average of three independent experiments is shown, with the standard deviation indicated on top of the histogram.

We conclude that state A and state B of the pre-40S particles here isolated with RIO1(kd)-StHA correspond to two very late maturation stages, just before and right after cleavage of the 18S rRNA 3’ end. These data indicate that rRNA cleavage and release of NOB1 and DIM2 coincide with the association of RIO1 in a stable conformation within the pre-40S particle and the incorporation of RPS26 in its final location. RIO1 enzymatic activity, and more specifically the release of the ATP hydrolysis products, seems required for its own dissociation from the particle rather than for the 18S-E pre-rRNA processing by NOB1.

### RPS26 is required for efficient rRNA cleavage and release of NOB1 and DIM2

States A and B of the RIO1(kd)-StHA pre-40S particles harbor two distinct conformations of the platform domain: in state A, the 18S-E pre-rRNA 3’ end is protected by DIM2, which prevents endonucleolytic cleavage by NOB1; in state B, the platform displays a mature conformation, with RPS26 replacing DIM2. Both RPS26 and RIO1 were shown to be necessary for processing of the 18S-E pre-rRNA to occur in human cells (O’Donohue et al., 2010; Widmann et al., 2012), which argues for the two proteins being involved in transition from state A to state B. Accommodation of RPS26 in its mature position requires prior release of DIM2, which occupies the binding site of RPS26.

In order to assess the role of RPS26 binding to pre-40S particles in 18S rRNA maturation as well as DIM2 and NOB1 release, we analyzed the RNA and protein composition of intermediate and late pre-40S particles purified from cells depleted of RPS26 using a specific siRNA (siRPS26). The level of 18S-E pre-rRNA cleavage in pre-40S particles was measured by RNase H analysis in pre-40S particles isolated using different baits (RIO1(kd)-StHA or RIO1(wt)-StHA, HASt-LTV1) (Figure 5, a and b). The amount of 18S-E pre-rRNA within early cytoplasmic pre-40S particles purified with HASt-LTV1 remained unchanged upon RPS26 depletion (Figure 5a), consistent with the absence of RPS26 in these particles (Ameismeier et al., 2018; Larburu et al., 2016; Wyler et al., 2011). In contrast, we observed a strong defect of 18S-E pre-rRNA processing in RIO1(kd)-StHA particles upon knockdown of RPS26, as evidenced by inversion of the mature to immature rRNA ratio when compared to the same particles isolated from cells treated with scramble siRNA. Similar observations were obtained when purifying pre-40S particles with a wild-type version of RIO1 as bait (RIO1(wt)-StHA pre-40S particles). These experiments indicate that absence of RPS26 impedes 18S-E pre-rRNA maturation, and that the action of RPS26 in 18S-E pre-rRNA cleavage takes place within RIO1-containing late cytoplasmic pre-40S particles.

**Figure 5.**
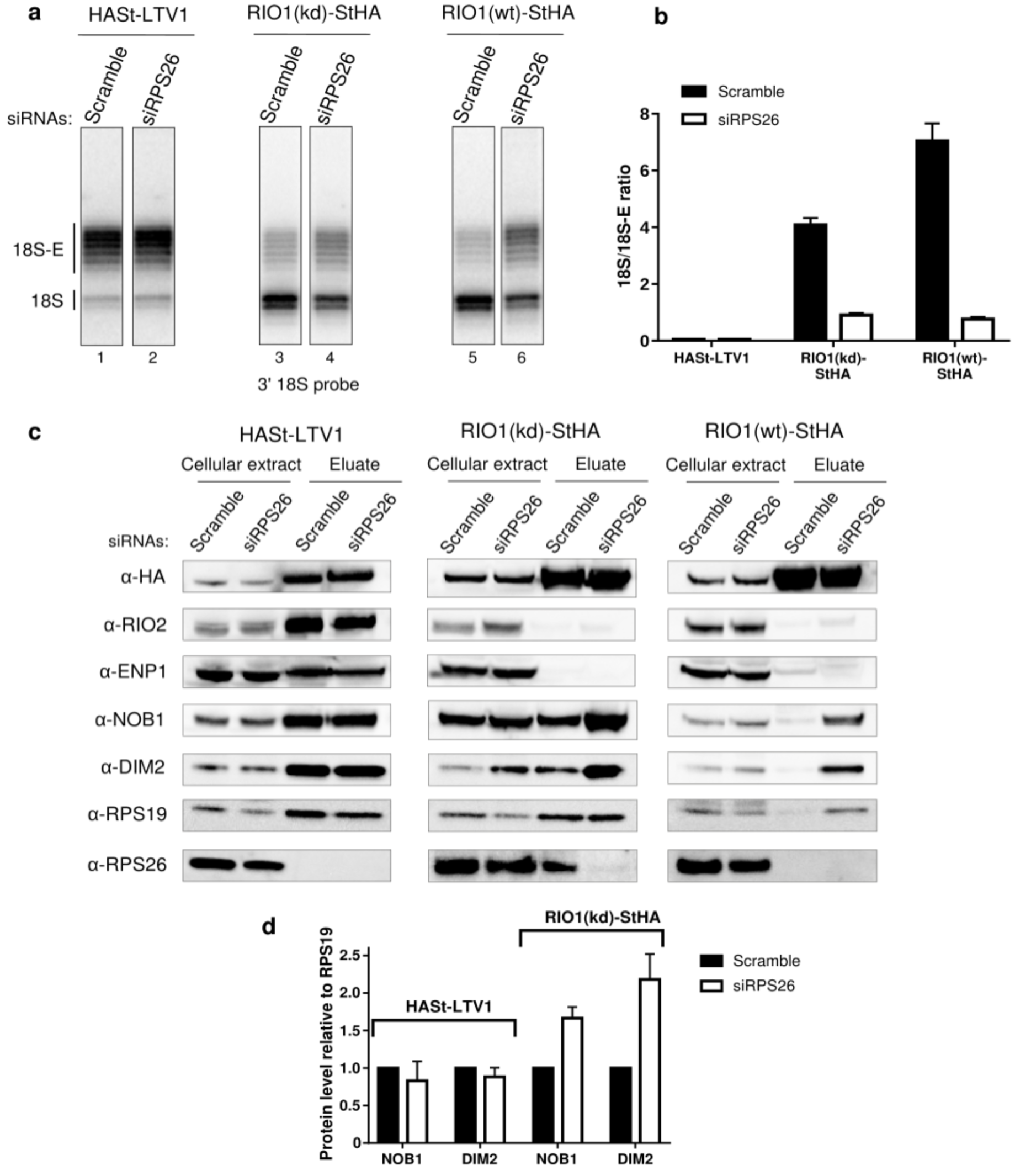
RPS26 is required for rRNA cleavage at site 3 as well as NOB1 and DIM2 release. HEK cell lines expressing tagged version of LTV1, the catalytically inactive RIO1-D324A (RIO1 (kd)) or wild-type RIO1 (RIO1(wt)) were treated with scramble or RPS26 siRNAs for 48 h. **a**, RNase H assays were conducted as in Figure 4 on rRNAs of pre-40S particles purified with the mentioned StHA tagged bait, either from RPS26-depleted cells or from control cells (scramble siRNA). **b**, Signals corresponding to the 18S-E and 18S rRNA detected in (a) were quantified and represented as the 18S/18S-E ratio for the different pre-40S particles. Error bars, s.d. (n=3) **c**, Cell extracts and purified particles were analysed by Western Blot using the indicated antibodies. **d**, Bands corresponding to DIM2 and NOB1 (in the eluates) were quantified, corrected for pre-40S particle loading (using RPS19) and normalized to the control condition (set to 1). Error bars, s.d. (n=3).

We then used Western blot analyses to monitor how RPS26 depletion influences the release of NOB1 and DIM2 (Figure 5c). As expected from previous studies (Larburu et al., 2016; Wyler et al., 2011), we did not detect RPS26 in early cytoplasmic HASt-LTV1 pre-40S particles. Consistently, knockdown of RPS26 did not influence the presence of NOB1 and DIM2 in these early cytoplasmic pre-40S particles, as attested by measuring the NOB1/RPS19 or DIM2/RPS19 ratios (Figure 5d), nor did it affect the dissociation of ENP1 and RIO2. In contrast, we observed a strong increase of NOB1 and DIM2 levels relative to RPS19 in RIO1(kd)-StHA pre-40S particles purified from RPS26 depleted cells when compared to control cells (Figure 5, c and d), which suggests that the absence of RPS26 prevents the release of NOB1 and DIM2 and traps the particles in state A. This hypothesis was further supported when we analyzed pre-40S particles purified using wild-type RIO1 as bait (RIO1(wt)-StHA). We have previously shown that RIO1(wt)-StHA does not co-purify efficiently with pre-40S particles when compared to the catalytically inactive version of RIO1 (Montellese et al., 2020; Widmann et al., 2012), which was confirmed here by the very low amount of both pre-40S RBFs and RPS19 detected by western blot (Figure 5c). However, upon RPS26 depletion, RIO1(wt)-StHA co-purified with late cytoplasmic pre-40S particles, as attested by the presence of NOB1 and DIM2 (Figure 5c), as well as that of 18S-E pre-rRNA (Figure 5a). These data indicate that RIO1 association to late cytoplasmic pre-40S particles is stabilized in the absence of RPS26, while release of NOB1 and DIM2 is inhibited. We conclude that RPS26 takes part in the mechanism triggering rRNA cleavage by NOB1 at site 3 as well as in dissociation of NOB1, DIM2 and RIO1.

### ATP binding by RIO1 stimulates rRNA cleavage by NOB1 in purified pre-40S particles

RIO protein kinases have been proposed to act as conformation-sensing ATPases rather than kinases (Ferreira-Cerca et al., 2014). Previous studies in the yeast *S. cerevisiae* have shown that rRNA cleavage by Nob1 in cytoplasmic pre-40S particles purified with tagged Rio1 was stimulated in vitro by ATP binding to Rio1 (Turowski et al., 2014). The D324A point mutation introduced in RIO1(kd)-StHA does not hamper fixation of ATP (Widmann et al., 2012), but the structure of RIO1(kd) in state B shows that the hydrolysis products are trapped in the catalytic site (Figure 3c). However, this defective catalytic activity does not block maturation of the 3’-end of 18S rRNA within RIO1(kd)-StHA pre-40S particles. This suggests that ATP binding to the human RIO1 catalytic site is sufficient for 18S-E pre-rRNA cleavage, similar to what was shown in yeast. In order to check whether ATP binding to RIO1 favors rRNA cleavage by NOB1, we purified RIO1(wt)-StHA and RIO1(kd)-StHA human pre-40S particles from RPS26 depleted cells to enrich pre-cleavage state A and performed in vitro cleavage assays.

Following the conditions established in yeast (Lebaron et al., 2012), we added Mn^2+^ either alone or supplemented with ATP or AMP-PNP (a non-hydrolysable analog of ATP) to the purified pre-40S particles, and monitored 18S rRNA 3’-end maturation using RNase H digestion. As shown in Figure 6, 18S-E pre-rRNA cleavage was stimulated, albeit with low efficiency, both by ATP and AMP-PNP in particles purified with RIO1(wt)-StHA or with RIO1(kd)-StHA. This result reinforces the idea that, like in yeast, ATP binding to RIO1 stimulates 18S-E cleavage at site 3 (Ferreira-Cerca et al., 2014; Turowski et al., 2014). Given the position of RIO1 in the vicinity of DIM2 and NOB1 revealed by the cryo-EM analysis, RIO1 might favor DIM2 displacement and subsequent activity of NOB1 through an ATP-driven local conformational change. Indeed, superimposition of state A and state B atomic models showed steric hindrance between the Cterminal regions of DIM2 and RIO1, which supports a mechanism of competition between these two proteins (Figure 6c). Nevertheless, the low yield of in vitro cleavage in these pre-40S particles, also reported in yeast, further supports the idea that RIO1 by itself is not sufficient to efficiently trigger processing by NOB1 in the absence of RPS26.

**Figure 6.**
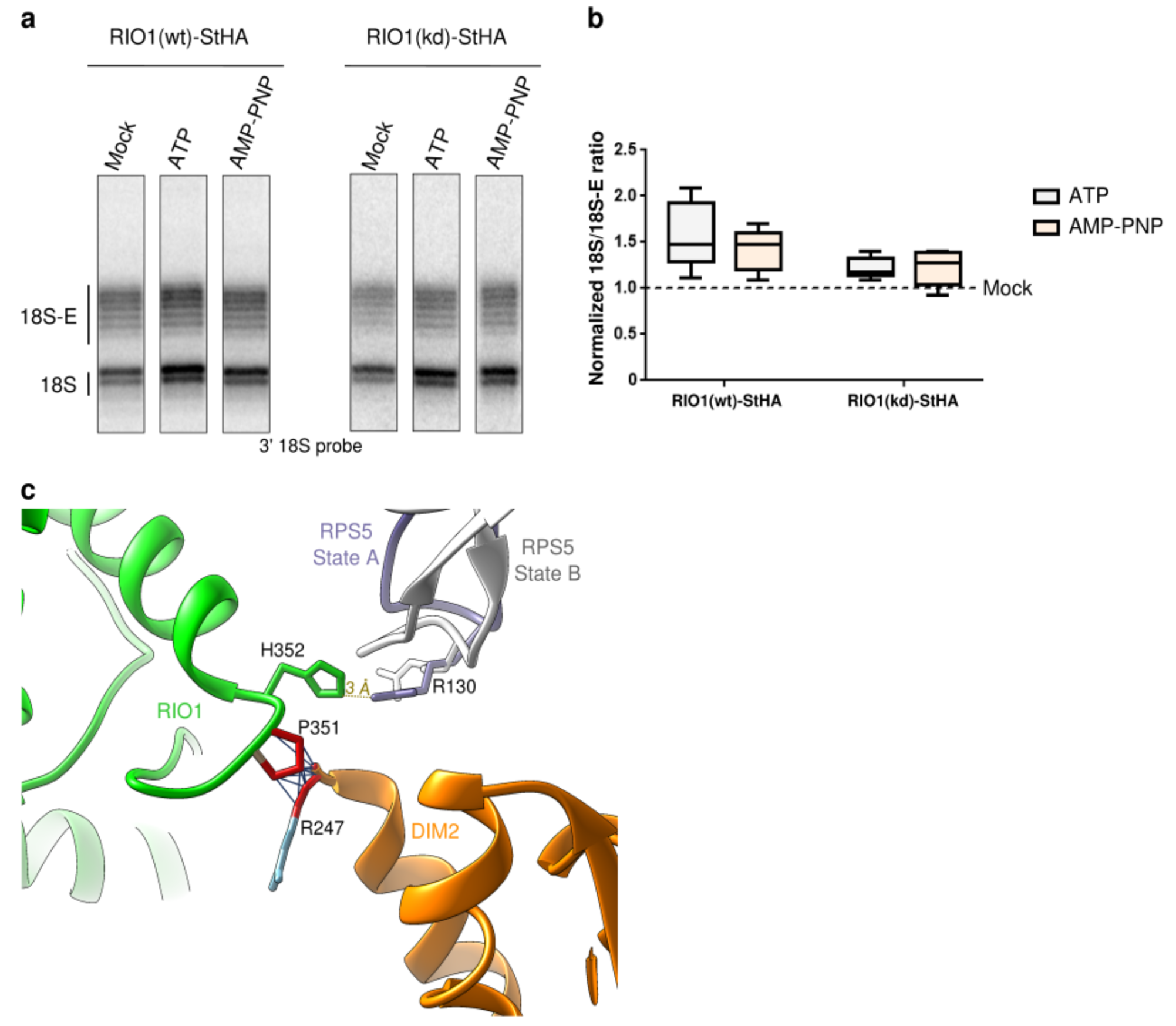
In vitro cleavage of the 18S-E pre-rRNA within pre-40S particles is stimulated by ATP addition. HEK cell lines expressing tagged versions of wild-type RIO1 (RIO1(wt)) or of the catalytically-inactive RIO1 (kd) were treated with scramble or RPS26 siRNAs for 48h to enrich particles in state A. Pre-40S particles were purified and incubated in the presence of 1 mM ATP, 1mM AMP-PNP, or without nucleotide (mock condition). **a**, RNAse H assays were performed on the RNAs extracted from the particles. **b**, The variation of cleavage efficiency with the different nucleotides is indicated by the 18S/18S-E ratio and normalized to the mock-treated sample (set to 1). The data correspond to five independent experiments. Analysis of the results with a unilateral paired Wilcoxon test (“sample greater than mock”) indicates p-values of 0.031 for samples RIO1(wt)-ATP, RIO1(wt)-AMP-PNP, RIO1(kd)-ATP, and 0.063 for RIO1(kd)-AMP-PNP. **c**, Superimposition of atomic models of State A and B reveals overlapping distances (grey lines) between atoms of Proline 351 from RIO1 (green) and of Arginine 247 from DIM2 (orange). RPS5, which seems to be repositioned upon association of RIO1 / dissociation of DIM2 from the pre-40S particle, is shown in violet (State A) or white (State B).

## DISCUSSION

A previous cryo-EM analysis of human pre-40S particles purified using DIM2 as bait revealed several successive structural states, in which the latest one carried NOB1, DIM2, TSR1 as well as a long alpha-helix attributed to LTV1 (Ameismeier et al., 2018). Here, by purification of a pre-40S particle via a catalytically inactive form of RIO1, we have uncovered two later structural states in which only the last RBFs found in pre-40S particles, namely NOB1, DIM2 and RIO1, were clearly detected. In addition, one of these states is posterior to 18S-E pre-rRNA cleavage at site 3, as indicated by detection of the mature 3’ end of the 18S rRNA, presence of RPS26, and absence of NOB1 and DIM2. These structures arguably correspond to the ultimate maturation stages of the pre-40S particles. Our results point towards a coordinated action of RIO1 and RPS26 in triggering the last 18S rRNA processing step by NOB1, following a sequence represented in Figure 7 and discussed below.

**Figure 7.**
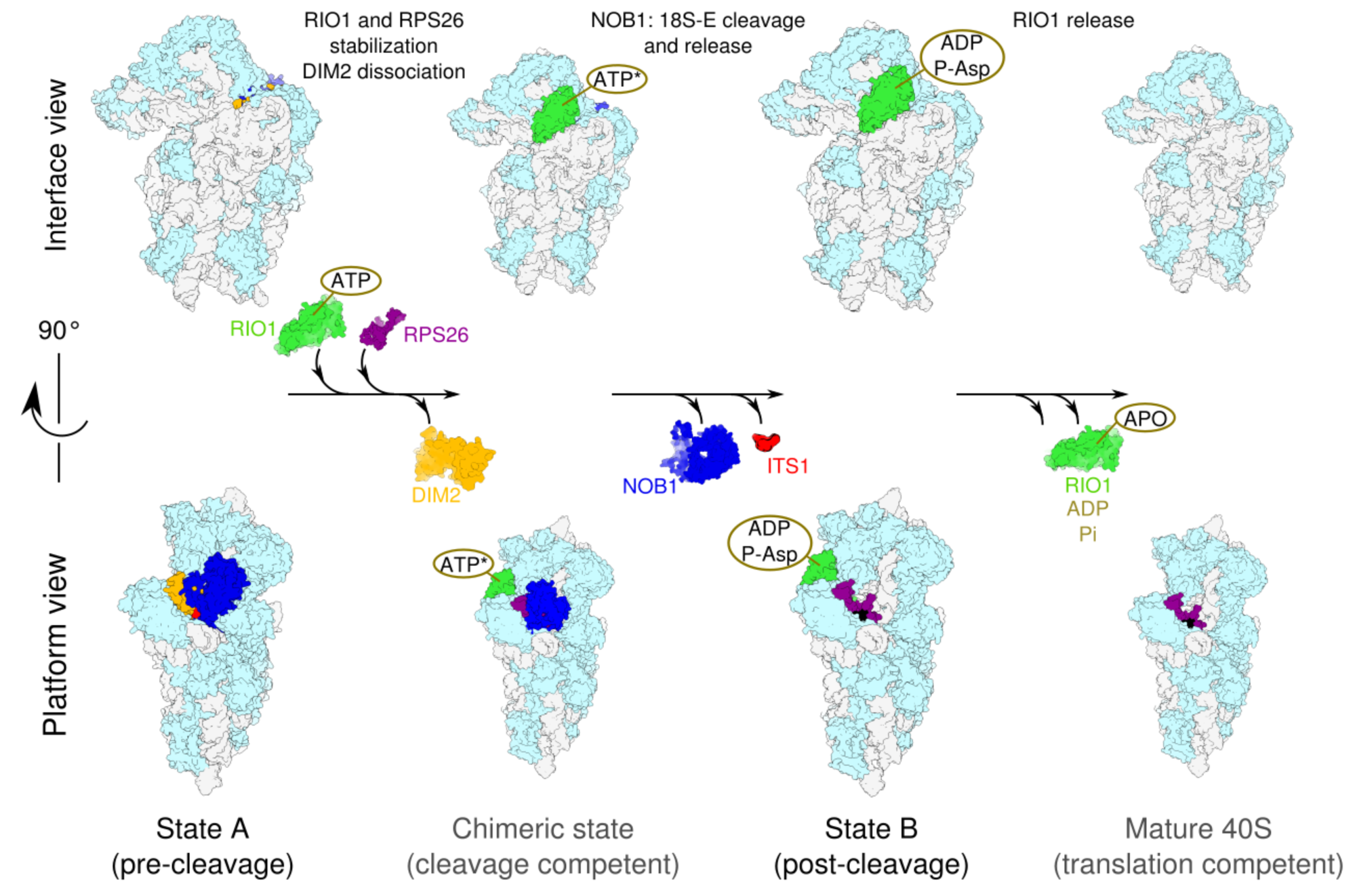
Model of the pre-40S last maturation steps triggered by RIO1 and RPS26. Upper and lower panels represent interface and platform views of the small ribosomal subunit, respectively. A putative ITS1 cleavage-competent state is shown to illustrate the transition between the pre-(State A) and post-cleavage (State B) structures which were resolved in this study. Status of ATP hydrolysis within RIO1 for this maturation state is not known, and thus marked as “ATP*”.

### The cooperative action of RIO1 and RPS26 unlocks 18S-E rRNA cleavage

Our results show that RIO1 binds to pre-40S particles in the same region as RIO2, which suggests that recruitment of RIO1 may simply follow the release of RIO2. The absence of a clear density for RIO1 in pre-cleavage state A suggests that RIO1 is initially loosely associated, while DIM2 and NOB1 are stably bound. In contrast, state B shows RIO1 in a stable conformation, while DIM2 and NOB1 are absent in this post-cleavage particle. Despite its overall impaired ATPase activity, the catalytic pocket of the RIO1-D324A mutant is occupied by ADP+pAsp341 (Figure 3c). This suggests that transition of pre-40S particles to state B involves ATP binding as well as the ATP hydrolysis activity of RIO-D324A, while trapping of ADP in the catalytic site prevents release from the matured particle. The structure of RIO1 was shown to switch to a so-called “active” form when complexed either with ATP or with its hydrolysis products ADP/pAsp (Ferreira-Cerca et al., 2014). ATP fixation in the catalytic pocket of RIO1 might trigger its stabilization on the head of the pre-40S particle and bring RIO1 to contact DIM2. Furthermore, our data reveal a possible steric clash between the C-terminal domains of RIO1 and DIM2 (Figure 6c). This suggests that RIO1 stabilization on pre-40S particles initiates DIM2 dislodging to uncover site 3 and allows rRNA cleavage by NOB1. Accordingly, our in vitro assays show that binding of ATP or AMP-PNP to RIO1 stimulates cleavage by NOB1 in human pre-40S particles (Figure 6), as also concluded before in yeast (Ferreira-Cerca et al., 2014). Also supporting this scenario, recent in vitro studies in yeast have shown that Rio1 forms a trimeric complex with Dim2 and Nob1 in the presence of AMP-PNP and not ADP (Parker et al., 2019). In addition, overexpression of Rio1 in yeast is sufficient to displace a catalytically-inactive form of Nob1 from pre-40S particles (Parker et al., 2019).

Nevertheless, RIO1 is not sufficient to efficiently trigger 18S-E rRNA cleavage together with DIM2 and NOB1 dissociation from pre-40S particles, as shown by the blocking of RIO1-StHA pre-40S particles in the pre-cleavage state upon RPS26 knockdown. RPS26 is the only component of the mature small ribosomal subunit missing in state A. Its binding site overlaps with that of DIM2 and its final positioning on the 40S subunits is thus directly dependent on the release of DIM2. While establishment of the full interaction of RIO1 with the head domain may provide a driving force to displace DIM2, RPS26 may potentiate its activity by competing with DIM2 binding. This suggests a simple model in which displacement of DIM2 upon RIO1 binding triggers the recruitment of RPS26 to the pre-40S particle, which in turn further displaces DIM2 from cleavage site 3. This cooperative action of RIO1 and RPS26 would then lead to a putative, probably short-lived cleavage-competent state (Figure 7), in which NOB1 can reach its substrate and process the 3-’end of the 18S rRNA. The status of ATP hydrolysis within RIO1 in this state is not known, and thus marked as “ATP*” in Figure 7. The release of ADP and dephosphorylation of phosphoaspartate pAsp341 would then trigger a conformational change within RIO1 allowing its dissociation from pre-40S particles, as proposed for yeast Rio1 as well as other RIO kinases family members (Ferreira-Cerca et al., 2012, 2014; Knüppel et al., 2018; Turowski et al., 2014). Along this view, RIO1 and RPS26 function as the two keys of a dual key lock that ensures a strict control of the ultimate steps of 40S subunit maturation before entry in translation.

### Human 40S subunit precursors do not form 80S-like particles unlike in yeast

In yeast, expression of the Rio1-D244A mutant (equivalent to human RIO1-D324A) was shown to promote the accumulation of 80S-like particles, with which it copurifies. In these 80S-like particles, pre-40S particles containing unprocessed 20S pre-rRNA, the last precursor to the 18S rRNA in yeast, are associated with large ribosomal subunits (Ferreira-Cerca et al., 2014; Turowski et al., 2014). Such particles were also observed with a number of other mutations targeting late-acting small ribosomal subunit biogenesis factors (Lebaron et al., 2012; Parker et al., 2019; Scaiola et al., 2018; Strunk et al., 2012) and were proposed to host the cleavage of the 20S pre-rRNA. Association of late pre-40S particles with large ribosomal subunits might serve as a checkpoint to verify that they are translation-competent after their final maturation (Lebaron et al., 2012; Strunk et al., 2012; Turowski et al., 2014). In addition, yeast 80S-like particles containing 20S pre-rRNA have been shown to be able to associate with polysomes and thus enter into the translating pool of ribosomes (Belhabich-Baumas et al., 2017; Parker et al., 2019; Soudet et al., 2010), suggesting some permissiveness between final pre-40S maturation and translation initiation events. In stark contrast, we found little evidence that human RIO1(kd)-StHA pre-ribosomal particles strongly associate to 60S subunits, neither by proteomics and Northern blot, nor by cryo-EM single particle analysis. Our results rather suggest that final rRNA maturation can occur within free pre-40S particles. Though ribosome biogenesis appears globally conserved within eukaryotic species, substantial differences in the pre-rRNA processing pathways, as well as in the dynamics of RBF association with pre-ribosomes in mammals and yeast have been demonstrated (for review, (Cerezo et al., 2019; Henras et al., 2015)). The absence of 80S-like particles in late human pre-40S particles might constitute a significant difference in the maturation pathway of human pre-ribosomal particles compared to yeast.

### RIO1 probes tRNA translocation capacities of pre-40S particles

Structural studies performed both in yeast and human showed that one of the roles of RIO2 was to prevent premature engagement of pre-40S particles in translation by physically obstructing the mRNA groove and tRNA interaction sites (Ameismeier et al., 2018; Heuer et al., 2017; Larburu et al., 2016; Scaiola et al., 2018; Strunk et al., 2012). In the post-cleavage state B, 18S rRNA G1639 appears to lock the RIO1(kd) catalytic pocket in its “active” conformation (Ferreira-Cerca et al., 2014) through a pi-stacking interaction with phenylalanine F328 (Figure 3c). Such an interaction does not seem to be established with RIO2, whose catalytic pocket is more distant from this guanosine (Ameismeier et al., 2018). The N7-methylated G1639 is universally conserved, and is involved in the correct positioning of tRNA in the P-site as well as its transfer to the E-site thanks to a switch mechanism (Malygin et al., 2013; Selmer et al., 2006). Thus, the presence of RIO1(kd) on the post-cleavage pre-40S particle prevents its entry into translation, by blocking rRNA-tRNA pairing. Through its cycle of association, ATPase hydrolysis and subsequent dissociation from pre-40S particles, RIO1 might thus act as a conformational and functional probe that assesses the tRNA translocation capacities of maturing rRNA. As suggested for other ribosome biogenesis factors, RIO1 would couple triggering of the last maturation step to functional proofreading of the ribosome.

## MATERIAL AND METHODS

### Cell culture and treatment with siRNA duplexes

HEK293 cells were cultured in Dulbecco’s modified Eagle’s medium (DMEM) supplemented with 10% fetal bovine serum and 1 mM sodium pyruvate. HEK293 FlpIn TRex cell lines expressing HASt-DIM2, HASt-LTV1, RIO1(wt)-StHA and RIO1(D324A)-StHA have been described previously (Wyler et al., 2011).

To knock down expression of the corresponding human protein genes, siRNA duplex of RPS26 (Eurogentec) (GenBank accession number: NM_001029): 5’-GGACAAGGCCAUUAAGAAA dTdT-3’) was added at a final concentration of 500 nM to 100 μl of cell suspension (50 × 10^6^ cells/ml diluted in Na phosphate buffer, pH 7.25, containing 250 mM sucrose and 1 mM MgCl_2_). After electrotransformation at 240 V, cells were plated and collected 48 h later. Control cells were electro-transformed with a scramble siRNA. Knockdown efficiency of siRNAs was assessed by quantitative PCR, using HPRT1 as an internal control.

### TAP purification of human pre-40S particles

To purify pre-ribosomal particles, the protocol described in (Wyler et al., 2011) was adapted as follows: expression of N-terminally HASt-tagged or C-terminally StHA-tagged bait proteins in stable HEK293 cells was induced with tetracycline (0,5μg/mL) 24 h prior to harvest. Cells were detached with PBS containing 0.5 mM EDTA and lysed in lysis buffer (10 mM Tris-HCl, pH 7.6, 100 mM KCl, 2 mM MgCl_2_, 1 mM DTT, 0.5% NP-40, supplemented with protease and phosphatase inhibitors) using a dounce homogenizer. Lysed cells were centrifuged (4500 g, 12 min) and the lysate was incubated with EZview Red Anti-HA Affinity Gel (Sigma aldrich) for 2 h in an overhead shaker. For electron microscopy studies, beads were washed six times with TAP buffer (20 mM Tris-HCl, pH 7.6, 100 mM KCl, 2 mM MgCl_2_, 1 mM DTT) and eluted by incubation in TAP buffer supplemented with 0.2 mg/mL HA peptide (Sigma aldrich). Eluates were washed and concentrated by Vivacon 2, 100 kDa MWCO centrifugal devices. For subsequent analyses by silver staining and Western blotting, beads were washed three times with TAP buffer and bound material was eluted with SDS sample buffer.

### Protein digestion and nanoLC-MS/MS analysis

100 μl of concentrated RIO1 pre-40S particles, corresponding to 3.5 μg of protein were reduced by incubation for 5 min at 95°C with 5 μL of Laemmli buffer containing 30 mM DTT, then alkylated with 90 mM iodoacetamide for 30 min at room temperature in the dark. The samples were then loaded and concentrated on an SDS-PAGE gel. For this purpose, the electrophoresis was stopped as soon as the proteins left the stacking gel to enter the resolving gel as one single band. The proteins, revealed with Instant Blue (Expedeon) for 20 minutes, were found in one blue band of around 5 mm width. The band was cut and washed before the in-gel digestion of the proteins overnight at 37°C with a solution of modified trypsin (sequence grade, Promega, Charbonnières, France). The resulting peptides were extracted from the gel by one round of incubation (15 min, 37°C) in 1% formic acid–acetonitrile (40%) and two rounds of incubation (15 min each, 37°C) in 1% formic acid–acetonitrile (1:1). The extracted fractions were air-dried. Tryptic peptides were resuspended in 14 μl of 2% acetonitrile and 0.05% trifluoroacetic acid (TFA). NanoLC-MS/MS analysis was performed in duplicate injections using an Ultimate 3000 nanoRS system (Dionex, Amsterdam, The Netherlands) coupled to an LTQ-Orbitrap Velos mass spectrometer (Thermo Fisher Scientific, Bremen, Germany) operating in positive mode. 5 μL of each sample were loaded onto a C18-precolumn (300 μm inner diameter x 5 mm) at 20 μL/min in 2% ACN, 0.05% TFA. After 5 min of desalting, the precolumn was switched online with the analytical C18 nanocolumn (75 μm inner diameter x 15 cm, packed in-house) equilibrated in 95 % solvent A (5 % ACN, 0.2% FA) and 5% solvent B (80% ACN, 0.2% FA). Peptides were eluted by using a 5-50% gradient of solvent B for 105 min, at a flow rate of 300 nL/min. The LTQ-Orbitrap Velos was operated in data-dependent acquisition mode with the XCalibur software. Survey scans MS were acquired in the Orbitrap, on the 300-2000 m/z (mass to charge ratio) range, with the resolution set to a value of 60,000 at m/z 400. Up to twenty of the most intense multiply charged ions (> 2+) per survey scan were selected for CID (collision-induced dissociation) fragmentation, and the resulting fragments were analyzed in the linear ion trap (LTQ). Dynamic exclusion was used within 60 s to prevent repetitive selection of the same peptide.

### Bioinformatic MS data analysis

Mascot (Mascot server v2.6.1; http://www.matrixscience.com) database search engine was used for peptide and protein identification using automatic decoy database search to calculate a false discovery rate (FDR). MS/MS spectra were compared to the SwissProt *H. sapiens* database (supplemented with the sequence of the D324A RIO1 mutant). Mass tolerance for MS and MS/MS was set at 5 ppm and 0.8 Da, respectively. The enzyme selectivity was set to full trypsin with two missed cleavages allowed. Protein modifications were fixed carbamidomethylation of cysteines, variable phosphorylation of serine, threonine and tyrosine, variable oxidation of methionine, variable acetylation of protein N-terminus. Proline software (Bouyssié et al., 2020) was used for the validation and the label-free quantification of identified proteins in each sample. Mascot identification results were imported into Proline. Search results were validated with a peptide rank=1 and at 1 % FDR both at PSM level (on Adjusted e-Value criterion) and protein sets level (on Modified Mudpit score criterion). Label-free quantification was performed for all proteins identified: peptides were quantified by extraction of MS signals in the corresponding raw files, and post-processing steps were applied to filter, normalize, and compute protein abundances. Peptide intensities were summarized in iBAQ values by dividing the protein abundances by the number of observable peptides (Schwanhäusser et al., 2011). Proteins with a ratio of observed peptide sequences over observable peptide sequences inferior to 30% were filtered out. Additionally, only the two most intense iBAQ logs (1E6 to 1E8) were represented in Figure 1 and Figure 1-figure supplement 2.

### Grid preparation and cryo-EM images acquisition

Cryo-EM grids were prepared and systematically checked at METI, Toulouse. Immediately after glow discharge, 3.5 μL of purified hRIO1(kd)-StHA particles (with RNA concentrations of ∼60ng/μL as estimated by Nanodrop measurement) were deposited onto QUANTIFOIL® holey carbon grids (R2/1, 300 mesh with a 2 nm continuous layer of carbon on top). Grids were plunge-frozen using a Leica EM-GP automat; temperature and humidity level of the loading chamber were maintained at 20°C and 95% respectively. Excess solution was blotted with a Whatman filter paper no.1 for 1.7-1.9 sec and grids were immediately plunged into liquid ethane (−183 °C).

Images were recorded on a Titan Krios electron microscope (FEI, ThermoFisher Scientific) located at EMBL, Heidelberg (Germany). The cryo-electron microscope was operating at 300 kV and was equipped with a Gatan K2 summit direct electron detector using counting mode. Automatic image acquisition was performed with SerialEM, at a magnification corresponding to a calibrated pixel size of 1.04 Å and a total electron dose of 29.88 e-/Å2 over 28 frames. Nominal defocus values ranged from −0.8 μm to −2.8 μm.

### Single particle analysis

9,494 stacks of frames were collected at EMBL. Frame stacks were aligned to correct for beam-induced motion using MotionCor2 (Zheng et al., 2017). Contrast Transfer Function (CTF) and defocus estimation was performed on the realigned stacks using CTFFIND4 (Rohou and Grigorieff, 2015). After selection upon CTF estimation quality, maximum resolution on their power spectra and visual checking, “good” micrographs were retained for further analysis. 2,126,610 particles were automatically picked, then extracted in boxes of 384×384 pixels, using the RELION 3.0 Autopick option. All subsequent image analysis was performed using RELION 3.0 (Zivanov et al., 2018) (Figure 2 – figure supplement 1). A first 2D classification was performed (on particles images binned by a factor of 8) to sort out ill-picked and methylosome particles. The 1,287,445 remaining particles were binned by a factor of 4 and subjected to a 3D classification in 5 classes, using the 40S subunit atomic model derived from the human ribosomal 3D structure (PDB-ID 6EK0) (Natchiar et al., 2018) low-pass filtered to 60 Å, as initial reference. One class harbored full 40S morphology and good level of details. The 484,429 particles from this class were re-extracted without imposing any binning factor, and a consensus 3D structure was obtained using RELION’s 3D auto-refine option, with an overall resolution of 2.9 Å for FSC=0.143 according to gold-standard FSC procedure. Particles were then submitted to focused 3D classifications with signal subtraction around the platform domain, to remove information coming from the body and head of the pre-40S particles, according to (Scheres, 2016). Two out of the five 3D classes yielded reconstructions with clear features. The first one, hereafter named state A contained 104,844 particles and was further auto-refined to 3.1 Å resolution according to gold-standard FSC procedure; the second one (state B) comprised 276,012 particles, and was auto-refined to 2.9 Å resolution. Subsequently, particles corresponding to the state A and state B “platform only” classes were retrieved from the dataset without signal subtraction, and submitted to auto-refinement, yielding maps of “full” state A and state B particles solved to 3.22 and 2.96 Å, respectively (Figure 2 – figure supplement 1, b and c). Because significant motion of the head and platform regions compared to the body were observed, multi-body refinement (Nakane et al., 2018) was performed for both states by dividing pre-40S particles in 3 main domains: body, head and platform. Multi-body refinement of state A gave rise to body, head and platform domains solved to 3.14, 3.17 and 3.30 Å, respectively after post-processing, while that of state B yielded resolutions of 2.98, 2.96 and 2.98 Å for the body, head and platform regions respectively (Figure 2 – figure supplement 1 c). Local resolution of all cryo-EM maps were estimated using the ResMap software (Kucukelbir et al., 2013) (Figure 2 – figure supplement 2).

### Interpretation of cryo-EM maps

Atomic models of pre-40S particle (PDB-ID 6G51) (Ameismeier et al., 2018) or the mature 40S subunit (PDB-ID 6EK0) (Natchiar et al., 2018) were first fitted in the cryo-EM maps of interest as rigid body using the “fit” command in UCSF Chimera (Pettersen et al., 2004). Body, head and platforms domains were modeled separately in the postprocessed multi-body “bodies” maps, and then adapted to the composite full maps to generate whole atomic models using UCSF Chimera and Coot (Emsley et al., 2010).

Final atomic models of state A (pre-cleavage) and B (post-cleavage) pre-40S particles were refined using REFMAC5 (Murshudov et al., 2011) and Phenix_RealSpace_Refine (Adams et al., 2010), with secondary structure restraints for proteins and RNA generated by ProSMART (Nicholls et al., 2014) and LIBG (Brown et al., 2015). Final model evaluation was done with MolProbity (Chen et al., 2010). Overfitting statistics were calculated by a random displacement of atoms in the model, followed by a refinement against one of the half-maps in REFMAC5, and Fourier shell correlation curves were calculated between the volume from the atomic model and each of the half-maps in REFMAC5 (Suppl. File 1).

Maps and models visualization was done with Coot and UCSF Chimera; figures were created using UCSF Chimera and ChimeraX (Goddard et al., 2018).

### RNase H digestion assay and RNA analysis

For RNase H digestion assays, 250 ng of pre-40S purified RNAs were denatured at 95°C for 6 min with a RNA/<u>DNA</u>/RNA reverse probe hybridizing in the 3’ end of 18S rRNA (probe RnaseH_3_Hyb1: 5’-UGUUACGACUUUU<u>ACTTCC</u>UCUAGAUAGUCAAGUUC-3’; 0,5 μl at 100 μM). After annealing by cooling down to room temperature for 20 min, the reaction mixture was diluted to 30 μl with a reaction mix containing 1 × RNase H reaction buffer, 25 μM DTT, 0.5 U/l RNasin (Promega) and 50 U RNase H (New England Biolabs), and incubated at 37°C for 30 min. The reaction was then blocked by addition of 0.3 M sodium acetate, pH 5.2 and 0.2 mM EDTA, and the RNAs were recovered by ethanol precipitation after phenol–chloroform–isoamylalcohol (25:24:1) extraction.

RNAs were then separated on a 12% polyacrylamide gel (19:1) in 1 × TBE buffer containing 7 M urea. RNAs were transferred to Hybond N+ nylon membrane (GE Healthcare, Orsay, France) and crosslinked under UV light. Membrane pre-hybridization was performed at 45°C in 6x SSC (saline-sodium citrate), 5x Denhardt’s solution, 0.5% SDS and 0.9 mg/ml tRNA. The 5’-radiolabeled oligonucleotide probe 3’18S (5’-GATCCTTCCGCAGGTTCACCTACG-3’) was added after 1 h and incubated overnight at 45°C. Membranes were washed twice for 10 min in 2x SSC and 0.1% SDS and once in 1x SSC and 0.1% SDS, and then exposed. Signals were acquired with a Typhoon Trio PhosphorImager and quantified using the ImageLab software.

The RNA ladder corresponds to a 131 nucleotides sequence comprizing the 3’18S probe and the T7 promoter sequence (Supplementary File 2). Using the oligonucleotides Lad_S and Lad_AS (see Supplementary File 2), the DNA template for the RNA ladder was produced by PCR. Then, RNA was synthesized by in vitro transcription of the PCR products using the T7 RNA polymerase (Promega kit). Then, alkaline hydrolysis was performed on 500ng of the synthesized RNA incubated in 50 mM sodium carbonate, pH 10.3, 1 mM EDTA for 5 min at 95°C.

### In vitro pre-ribosome maturation assay

The method of pre-40S particles purification above described was followed and instead of elution, beads were then washed 3 times with 1 ml of TAP buffer. Most of the supernatant was discarded and 50 μl of buffer X (50 mM Tris-HCl pH 7.5, 150 mM NaCl, 5 mM MnCl_2_, 1 mM DTT, 1% Triton and 10% glycerol) added to the beads (recovered by resuspension in 20 μl of remaining buffer 2) to reach a final volume of 70 μl. Nucleotides were added when required at a final concentration of 1 mM. Reactions were incubated at 37°C for 1 h and RNAs were then immediately extracted with Tri Reagent.

## DATA AVAILABILITY

Mass spectrometry proteomics data have been deposited to the ProteomeXchange Consortium via the PRIDE partner repository with the dataset identifier PXD019270. Cryo-EM maps have been deposited in the Electron Microscopy Data Bank (EMDB), under the accession codes : EMD-11440 (State A multi-body composite map); EMD-11441 (State B multi-body composite map); EMD-11446 (State A, head); EMD-11445 (State A, body); EMD-11447 (State A, platform); EMD-11443 (State B, head); EMD-11442 (State B, body); EMD-11444 (State B, platform). Atomic coordinate models of State A and State B RIO1(kd)-StHA pre-40S particles have been deposited in the Protein Data Bank (PDB), with respective PDB accession codes 6ZUO and 6ZV6.

## ACKNOWLEDGEMENTS

This work was funded by the Agence Nationale de la Recherche (ANR 16-CE11-0029), the CNRS, the University of Toulouse-Paul Sabatier, the Région Occitanie, the European Union (Fonds Européens de Développement Régional, FEDER) and Toulouse Métropole. UK received funding by the Swiss National Science Foundation (grant 31003A_166565 and the NCCR ‘RNA and disease’). Cryo-EM image acquisition was performed on the METI facility (CBI, Toulouse) and at the EMBL in Heidelberg with financial support from the iNext European initiative. Cryo-EM image analysis was performed using HPC resources from CALMIP (Grant 2018-[P1406]). We would like to thank the engineers and staff working on these facilities for their great help, as well as Marion Aguirrebengoa for her help on statistical analysis and Odile Burlet-Schiltz for her support.

## SUPPLEMENTARY MATERIAL

**Figure 1 – figure supplement 1.**
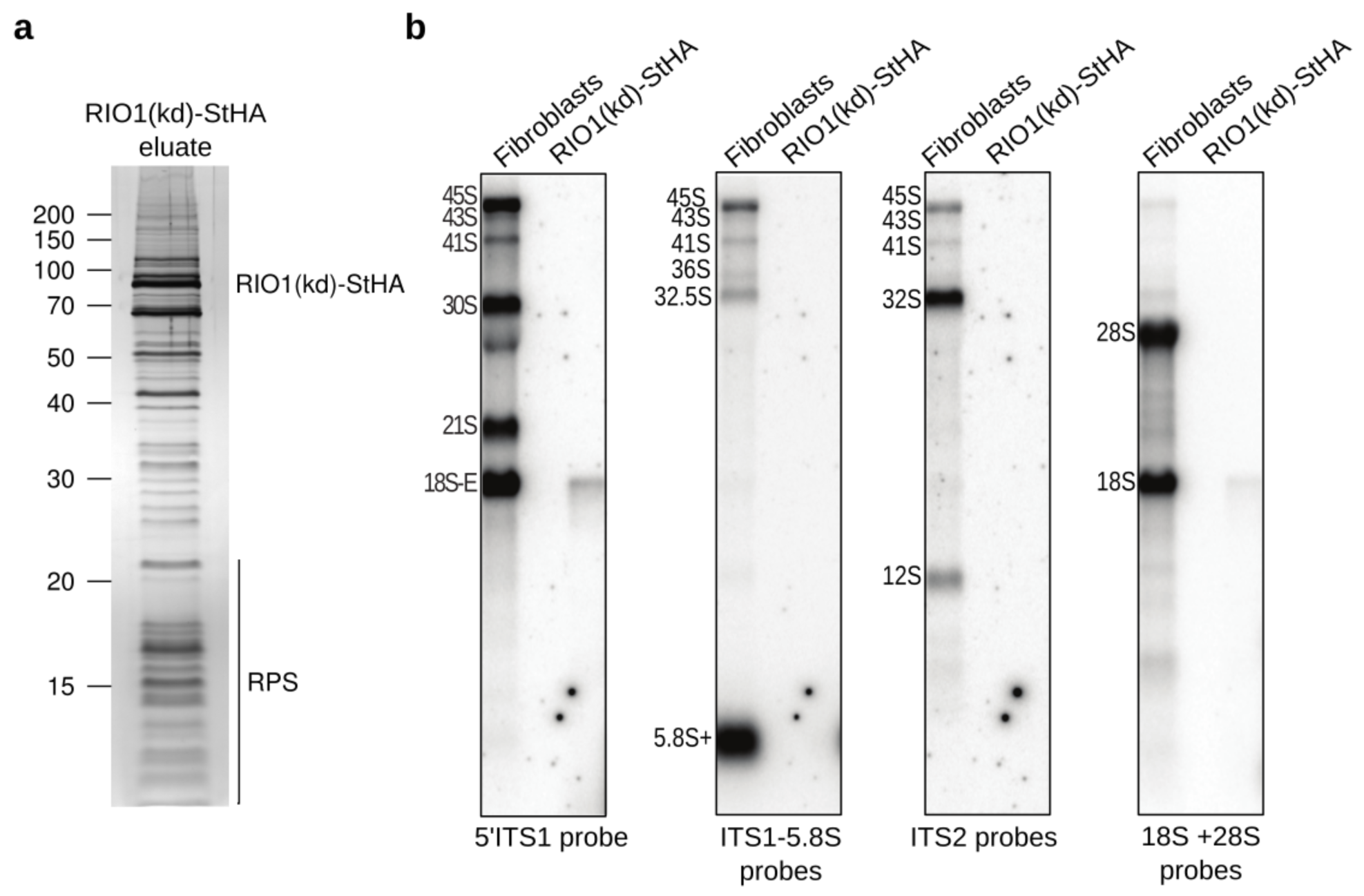
Tandem affinity purification of RIO1(kd)-StHA-containing particles. **a**, RIO1(kd)-StHA purification products analyzed by SDS-PAGE followed by silver staining. **b**, Northern blot analysis of the RNA content of the RIO1(kd)-StHA pre-40S particles (right lanes) compared to fibroblast total RNA (left lanes), revealed with 5’ITS1, ITS1-5.8S, ITS2 and 18S+28S probes (0.2 μg / lane).

**Figure 1 – figure supplement 2.**
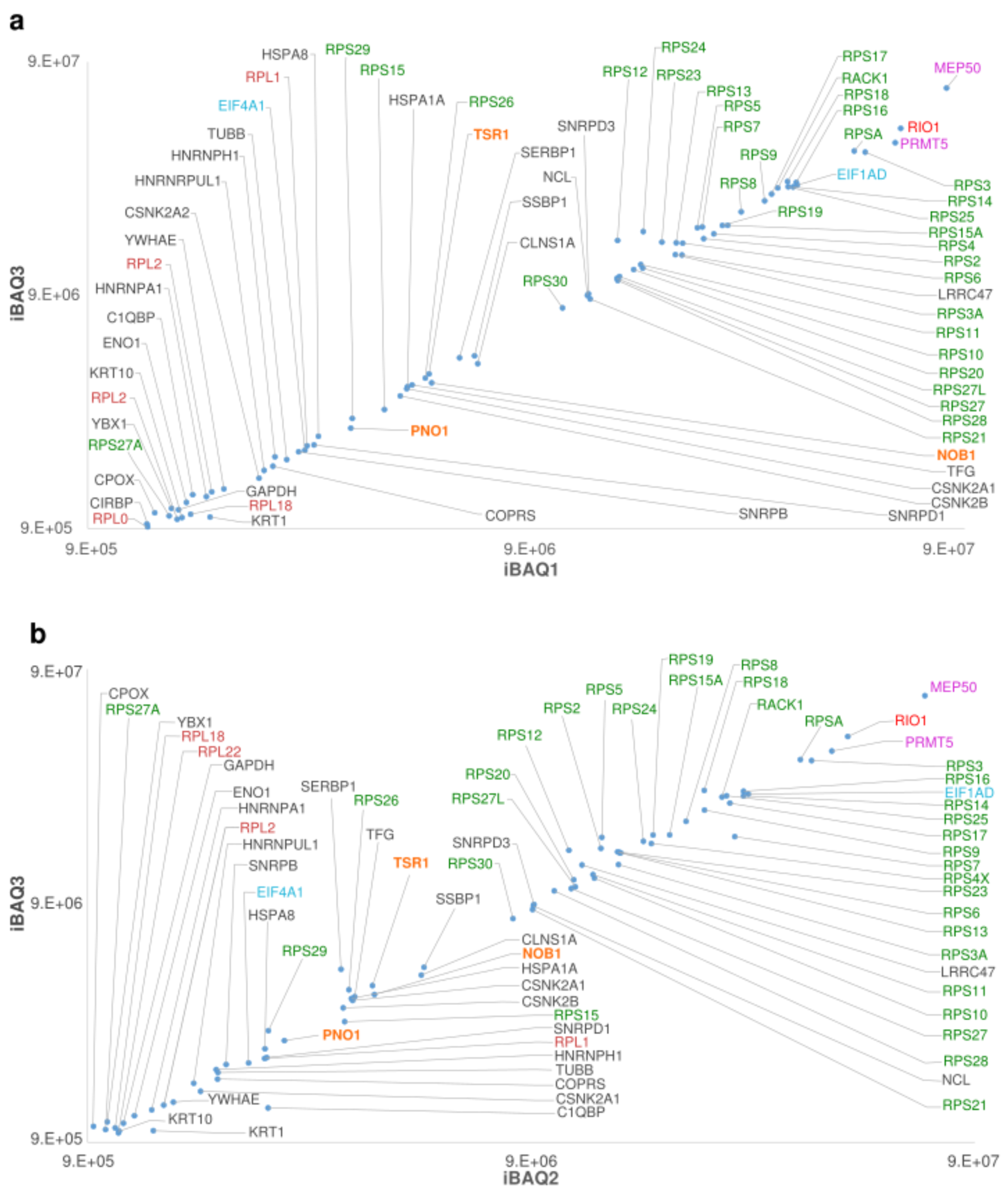
Label-free bottom-up proteomic analyses of RIO1(kd)-StHA co-purified proteins. All plotted proteins harbor an observed/observable peptide ratio > 30%; names of proteins with the 75 highest ratios are displayed, with color codes as indicated on the graph. **a**, iBAQs of experimental replicate 1 (iBAQ1) against experimental replicate 3 (iBAQ3). **b**, iBAQs of experimental replicate 2 (iBAQ2) against experimental replicate 3 (iBAQ3).

**Figure 2 – figure supplement 1.**
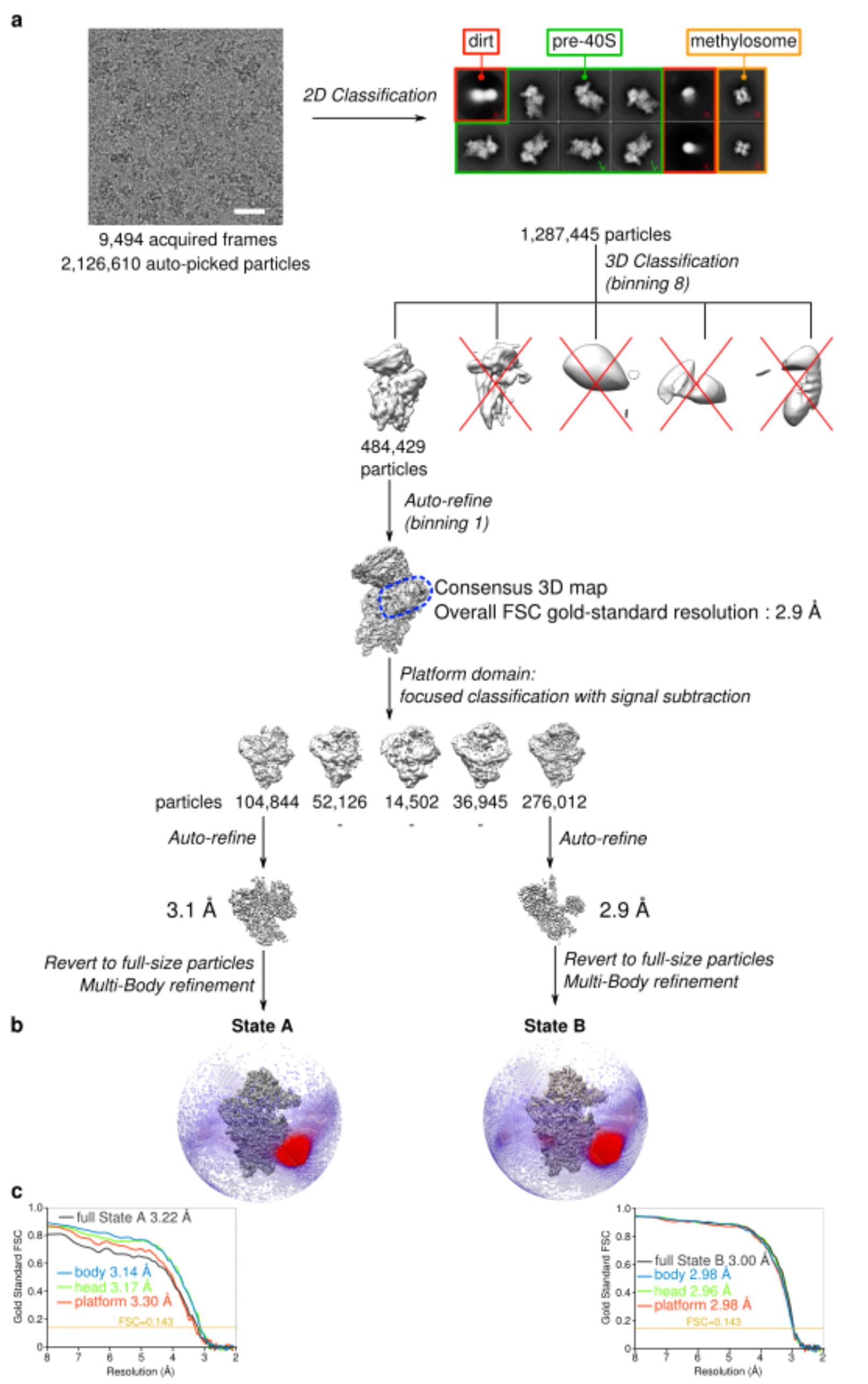
Cryo-EM image processing scheme. **a**, Single particle analysis strategy applied for obtaining the states A and B RIO1(kd)-StHA structures. **b**, State A (left panel) and B (right panels) cryo-EM maps and Euler angle distribution as seen along the interface side; **c**, Gold standard FSC curves for the various cryo-EM maps obtained for state A (left panel) and B (right panel).

**Figure 2 – figure supplement 2.**
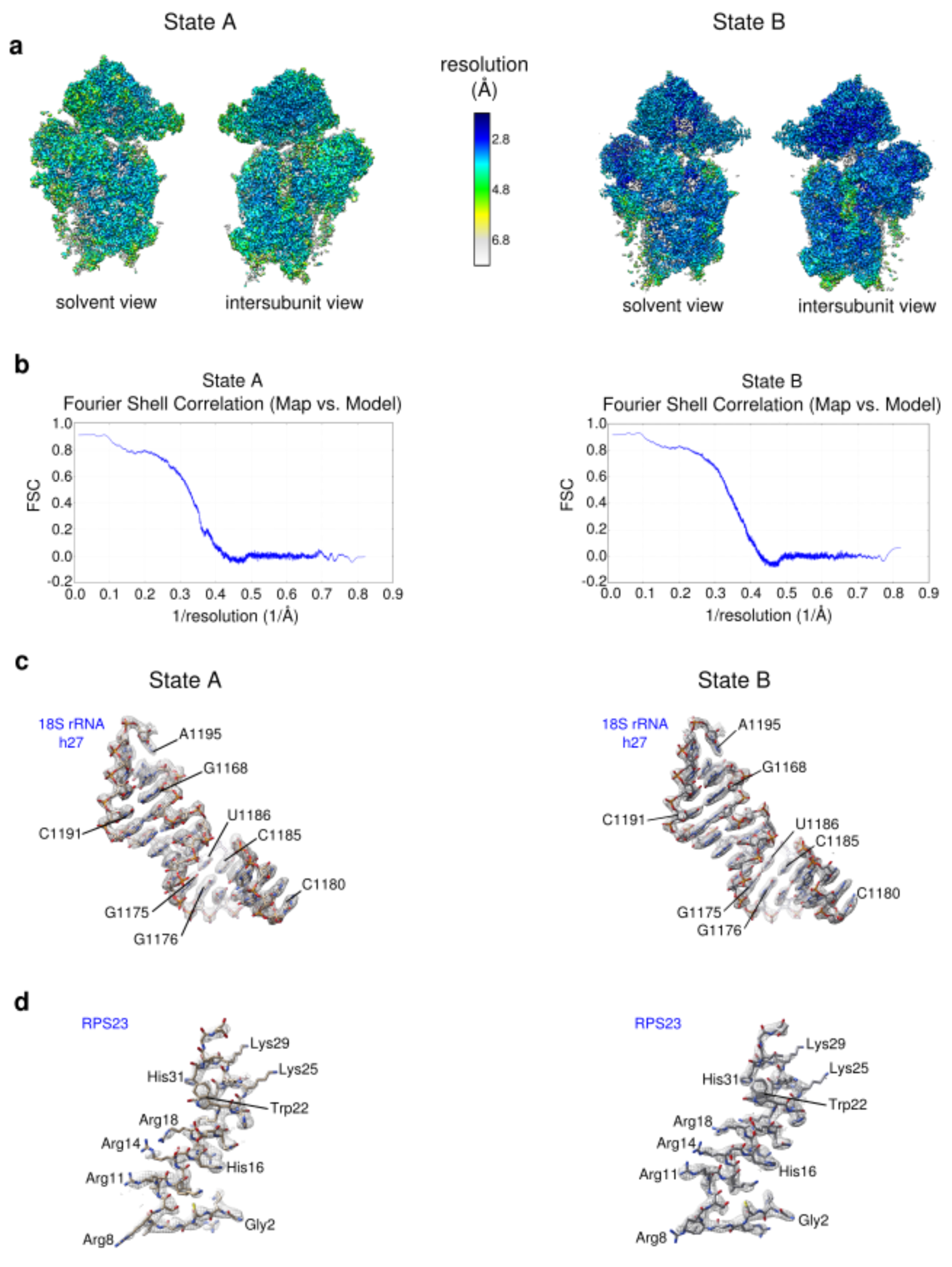
Details of the cryo-EM structures / model validation. **a**, Composite post-processed maps of state A (left panel) and state B (right panel) colored according to local resolution estimations calculated by the Resmap program (Kucukelbir et al., 2013). **b**, Validation of the atomic models derived from the cryo-EM maps of hRIO1(kd)-StHA pre-40S particles: model to map Fourier Shell Correlation estimation calculated by Molprobity for state A (left panel) and state B cryo-EM data. **c, d**, Details of the atomic models of state A (left panel) and state B pre-40S particles showing 18S rRNA helix 27 (c) and N-terminal domain of RPS23 (d); cryo-EM map density is represented by a grey mesh.

**Figure 3 – figure supplement 1.**
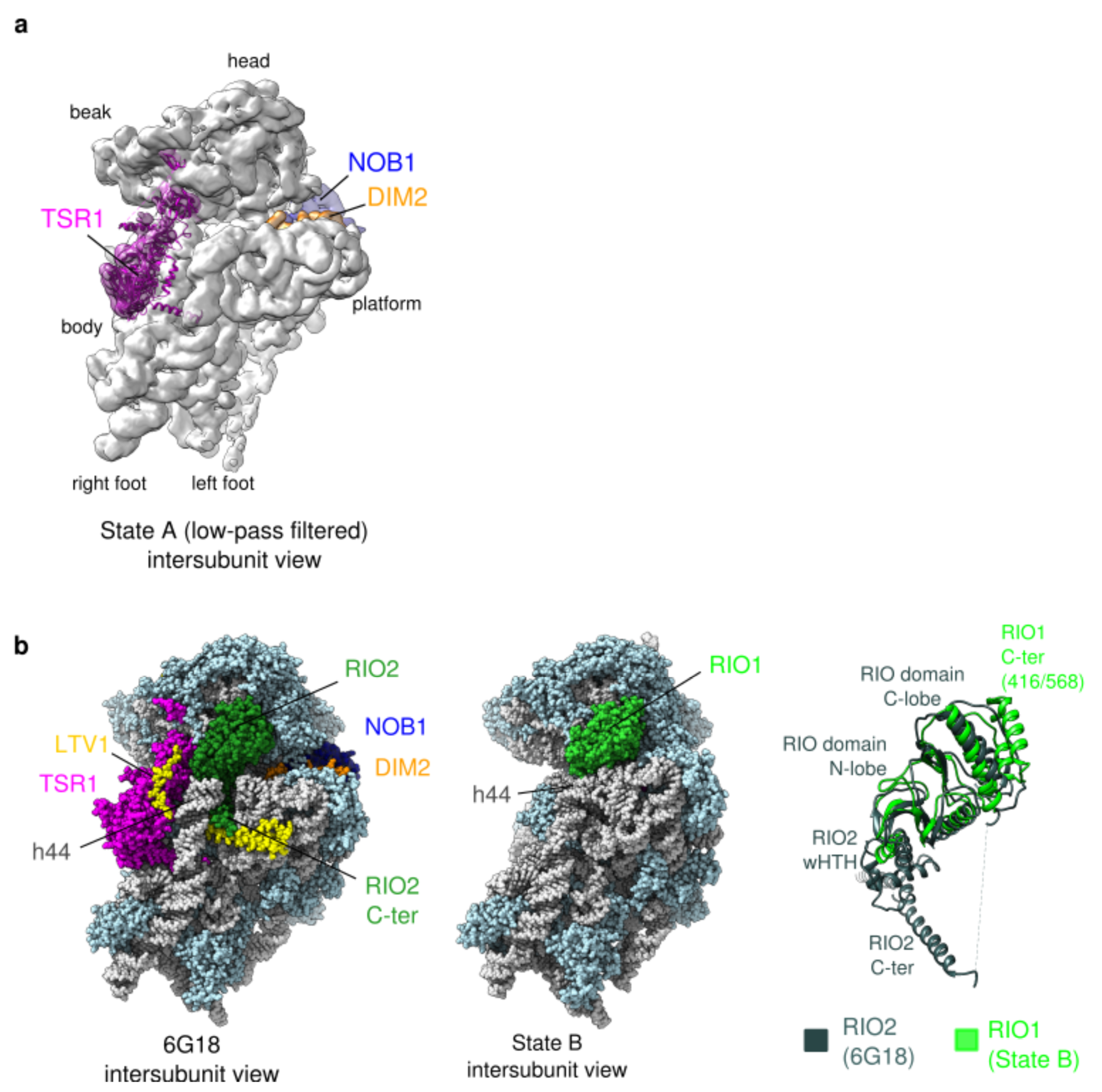
Details of the structural analysis of hRIO1(kd)-StHA pre-40S particles. **a**, Low-pass filtered cryo-EM map of state A reveals a density on the intersubunit side under the beak (colored in magenta) which can unambiguously accommodate the atomic structure of TSR1 (ribbons colored in magenta; atomic model from ((Ameismeier et al., 2018), PDB ID: 6G18). **b**, RIO1 replaces RIO2 on the back of the head of pre-40S particles. Left panel, atomic model of intermediate cytoplasmic pre-40S particle (state C of (Ameismeier et al., 2018), PDB ID: 6G18) featuring RIO2, displayed in dark green. The upper part of rRNA helix h44 is lifted from the rest of the body, allowing the C-terminal domain of RIO2 as well as a domain of LTV1 (yellow) to be deeply inserted within the pre-40S particle. Middle panel, atomic model of state B of hRIO1(kd)-StHA pre-40S particles. RIO1 is shown in light green. The upper part of rRNA helix h44 is close to the rest of the particle, preventing the C-terminal domain of RIO1 to occupy this region. Right panel, superimposition of RIO2 (dark green) and RIO1 as found on the two pre-40S particles herein presented.

**Figure 4 – figure supplement 1.**
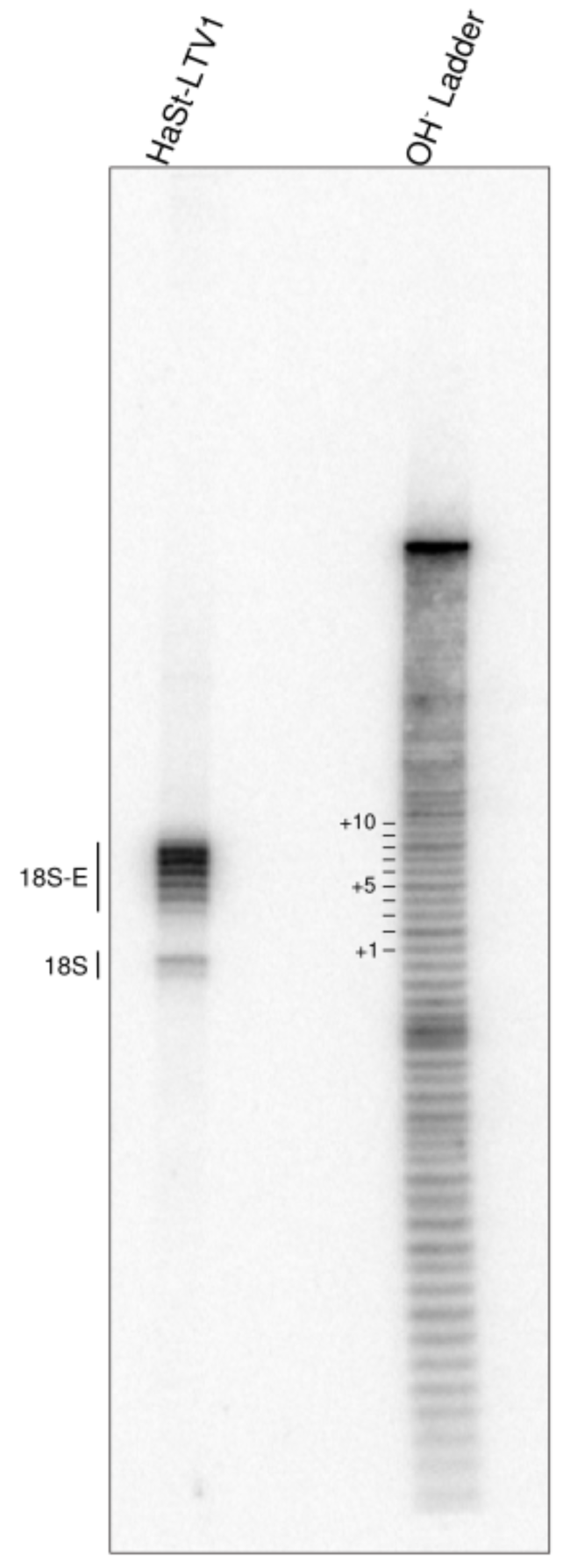
Assessment of the size of the ITS1 in purified pre-40S particles. Comparison of RNAse H digestion and alkaline hydrolysis assays shows nucleotide resolution between bands. Left panel, RNase H digestion of rRNAs from pre-40S particles purified using a HASt tagged version of LTV1 as bait (HASt-LTV1). Right panel, alkaline hydrolysis (OH^-^ ladder) of an RNA molecule containing the 18S-ITS1 sequence recognized by the 3’18S probe at its 5’ end (see Suppl. File 2). The samples were fractionated ob a 12% polyacrylamide gel and northern blot was probed with the 3’18S probe.

**Supplementary File 1.**
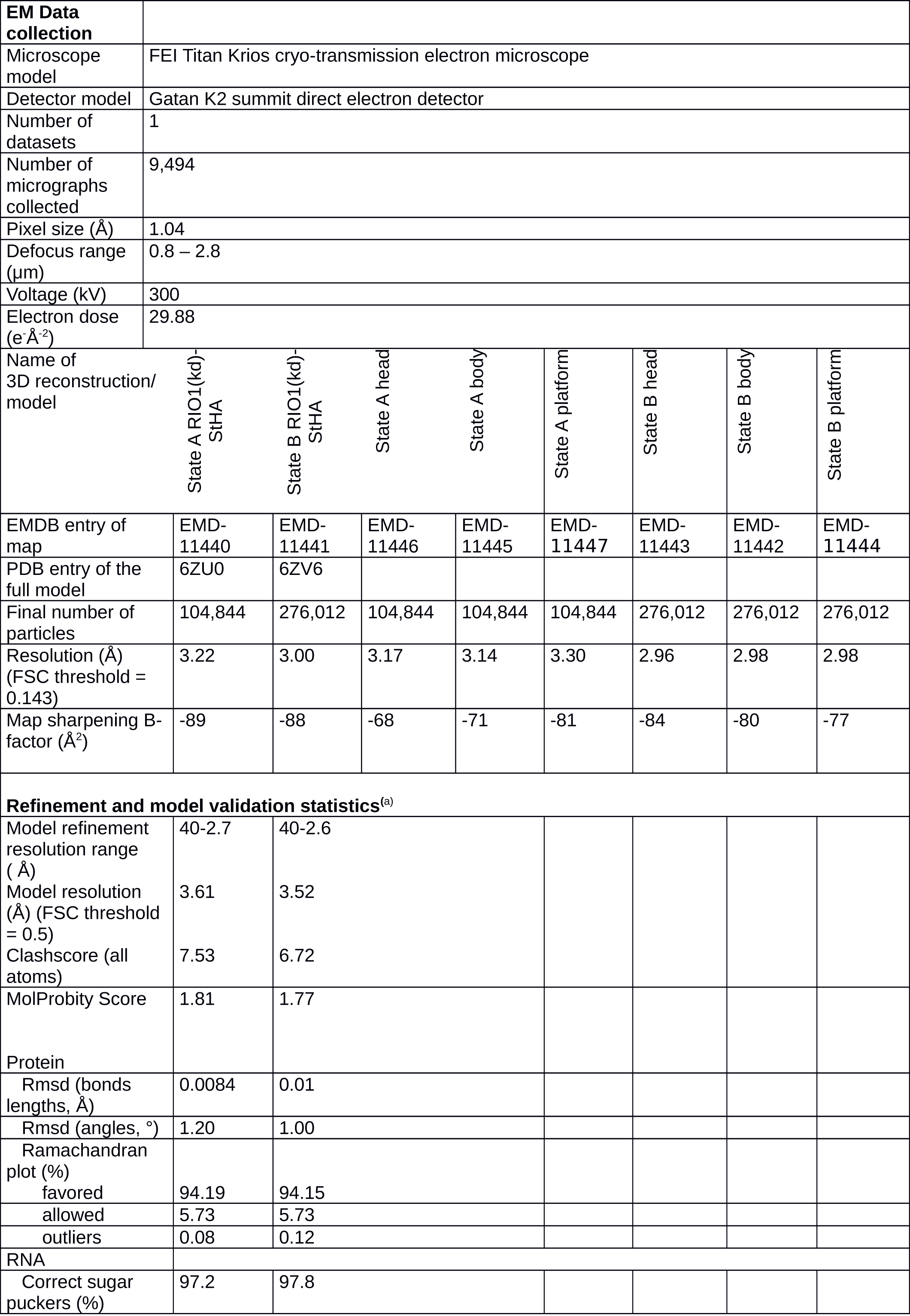
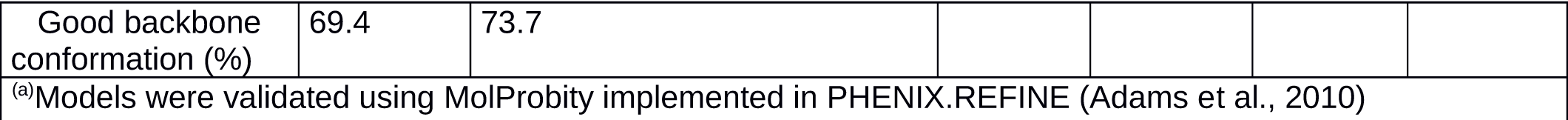
Cryo-EM data collection, atomic models refinement and validation statistics.

|  |  |  |  |  |  |  |  |  |
| --- | --- | --- | --- | --- | --- | --- | --- | --- |
| **EM Data collection** |  |  |  |  |  |  |  |  |
| Microscope model | FEI Titan Krios cryo-transmission electron microscope |  |  |  |  |  |  |  |
| Detector model | Gatan K2 summit direct electron detector |  |  |  |  |  |  |  |
| Number of datasets | 1 |  |  |  |  |  |  |  |
| Number of micrographs collected | 9,494 |  |  |  |  |  |  |  |
| Pixel size (Å) | 1.04 |  |  |  |  |  |  |  |
| Defocus range (µm) | 0.8 – 2.8 |  |  |  |  |  |  |  |
| Voltage (kV) | 300 |  |  |  |  |  |  |  |
| Electron dose (e<sup>-</sup>Å<sup>-2</sup>) | 29.88 |  |  |  |  |  |  |  |
| Name of 3D reconstruction/ model | State A RIO1(kd)-StHA | State B RIO1(kd)-StHA | State A head | State A body | State A platform | State B head | State B body | State B platform |
| EMDB entry of map | EMD-11440 | EMD-11441 | EMD-11446 | EMD-11445 | EMD-11447 | EMD-11443 | EMD-11442 | EMD-11444 |
| PDB entry of the full model | 6ZU0 | 6ZV6 |  |  |  |  |  |  |
| Final number of particles | 104,844 | 276,012 | 104,844 | 104,844 | 104,844 | 276,012 | 276,012 | 276,012 |
| Resolution (Å) (FSC threshold = 0.143) | 3.22 | 3.00 | 3.17 | 3.14 | 3.30 | 2.96 | 2.98 | 2.98 |
| Map sharpening B-factor (Å<sup>2</sup>) | -89 | -88 | -68 | -71 | -81 | -84 | -80 | -77 |
| **Refinement and model validation statistics<sup>(a)</sup>** |  |  |  |  |  |  |  |  |
| Model refinement resolution range ( Å) | 40-2.7 | 40-2.6 |  |  |  |  |  |  |
| Model resolution (Å) (FSC threshold = 0.5) | 3.61 | 3.52 |  |  |  |  |  |  |
| Clashscore (all atoms) | 7.53 | 6.72 |  |  |  |  |  |  |
| MolProbity Score | 1.81 | 1.77 |  |  |  |  |  |  |
| Protein |  |  |  |  |  |  |  |  |
| Rmsd (bonds lengths, Å) | 0.0084 | 0.01 |  |  |  |  |  |  |
| Rmsd (angles, °) | 1.20 | 1.00 |  |  |  |  |  |  |
| Ramachandran plot (%) |  |  |  |  |  |  |  |  |
| favored | 94.19 | 94.15 |  |  |  |  |  |  |
| allowed | 5.73 | 5.73 |  |  |  |  |  |  |
| outliers | 0.08 | 0.12 |  |  |  |  |  |  |
| RNA |  |  |  |  |  |  |  |  |
| Correct sugar puckers (%) | 97.2 | 97.8 |  |  |  |  |  |  |

|  |  |  |
| --- | --- | --- |
| Good backbone conformation (%) | 69.4 | 73.7 |

<sup>(a)</sup>Models were validated using MolProbity implemented in PHENIX.REFINE (Adams et al., 2010)

**Supplementary File 2.**
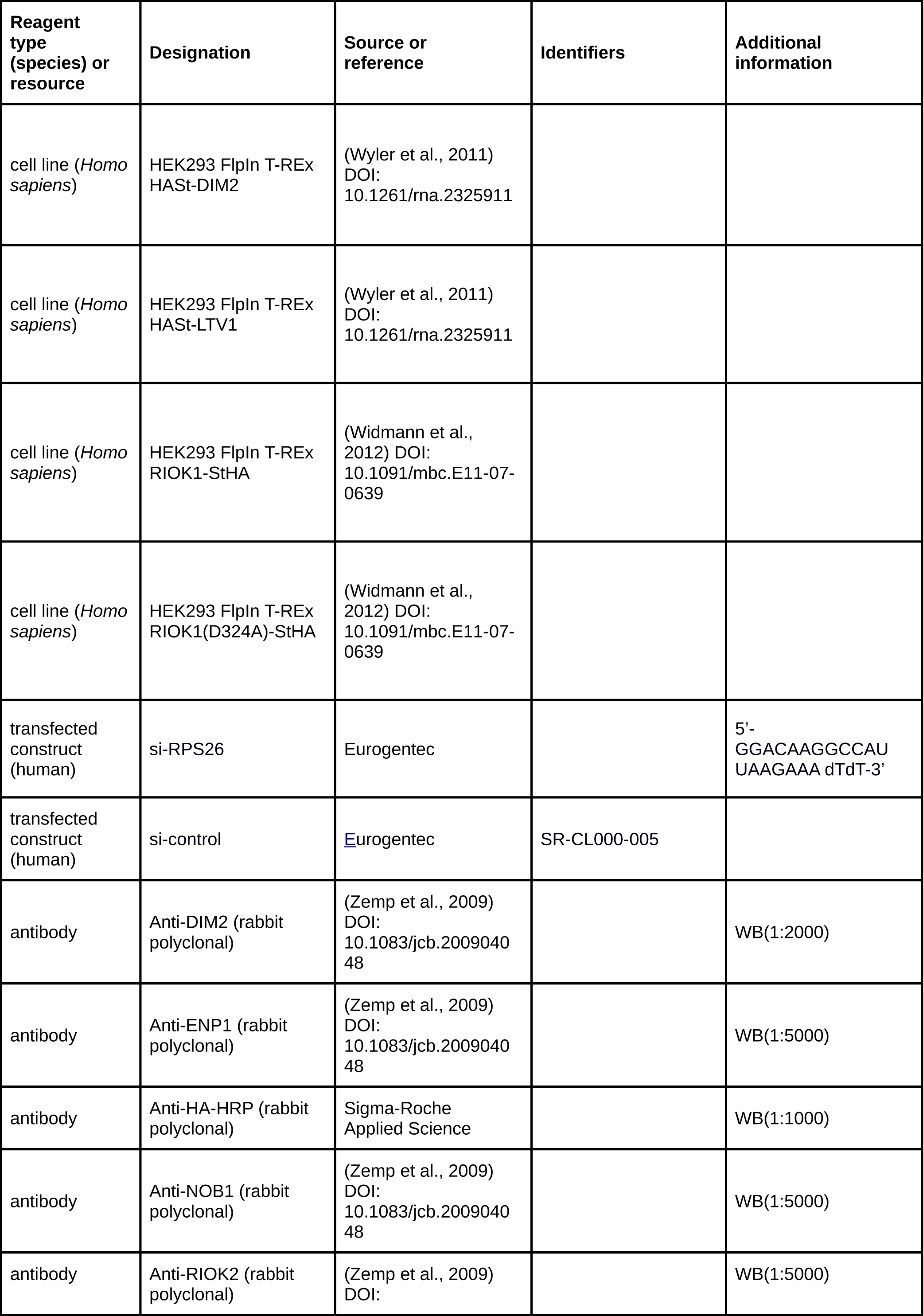
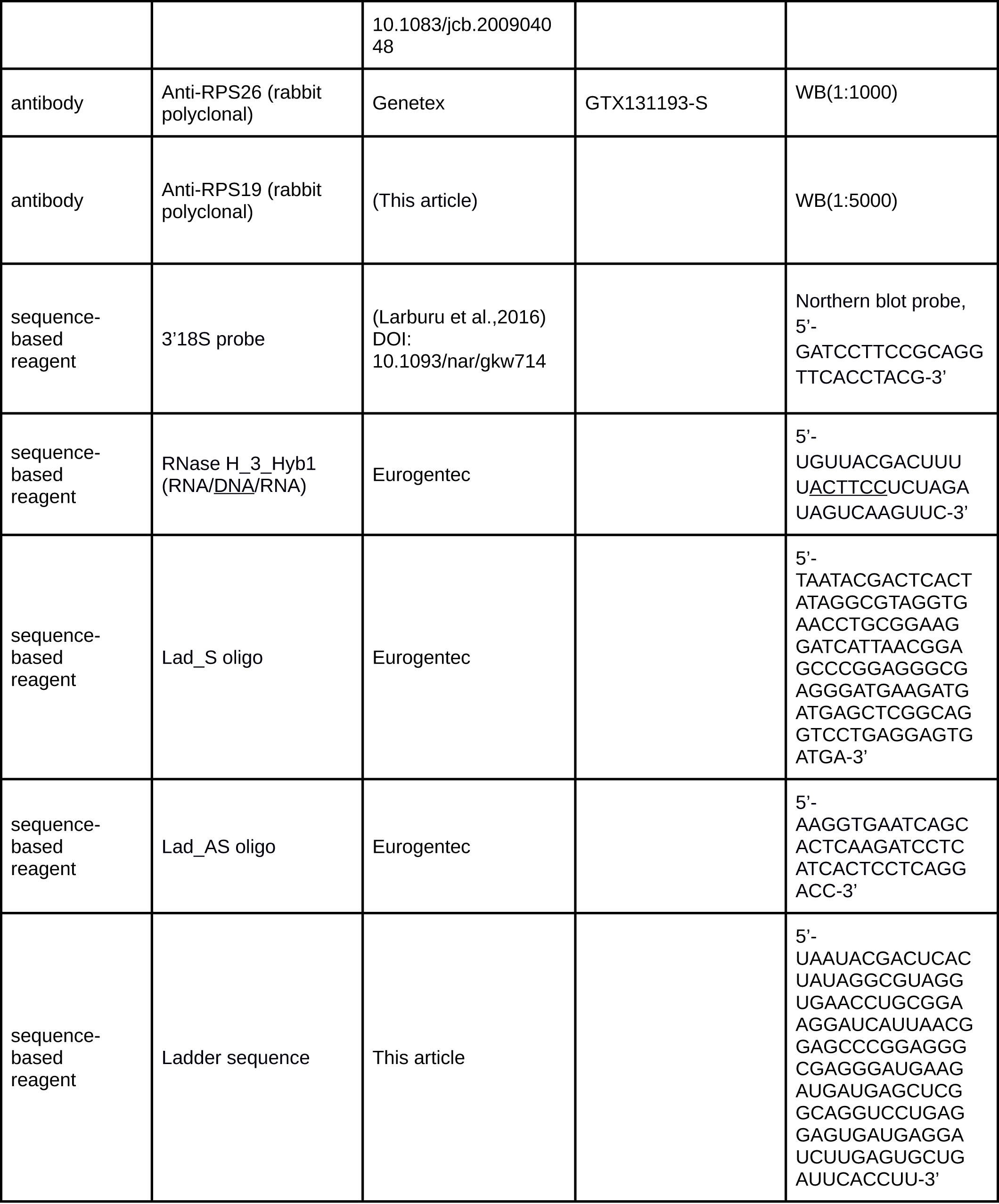
Key Resources Table

